# Phyloformer: Fast, accurate and versatile phylogenetic reconstruction with deep neural networks

**DOI:** 10.1101/2024.06.17.599404

**Authors:** Luca Nesterenko, Luc Blassel, Philippe Veber, Bastien Boussau, Laurent Jacob

## Abstract

Phylogenetic inference aims at reconstructing the tree describing the evolution of a set of sequences descending from a common ancestor. The high computational cost of state-of-the-art Maximum likelihood and Bayesian inference methods limits their usability under realistic evolutionary models. Harnessing recent advances in likelihood-free inference and geometric deep learning, we introduce Phyloformer, a fast and accurate method for evolutionary distance estimation and phylogenetic reconstruction. Sampling many trees and sequences under an evolutionary model, we train the network to learn a function that enables predicting the former from the latter. Under a commonly used model of protein sequence evolution and exploiting GPU acceleration, it outpaces fast distance methods while matching maximum likelihood accuracy on simulated and empirical data. Under more complex models, some of which include dependencies between sites, it outperforms other methods. Our results pave the way for the adoption of sophisticated realistic models for phylogenetic inference.

## 1 Introduction

Phylogenetics, the reconstruction of evolutionary relationships between biological entities, is used in many research domains to provide essential insights into evolutionary processes. It is employed in epidemiology to track viral spread [1], in virology to identify events of recombination [2], in biochemistry to evaluate functional constraints operating on sequences [3], in ecology to characterize biodiversity [4]. Central to these works, the phylogeny is a binary tree whose internal nodes correspond to ancestral entities, branches represent the amount of evolutionary divergence, and leaves correspond to extant entities. Most of the time, molecular phylogenies are estimated from aligned nucleotide or amino acid sequences using probabilistic model-based approaches in the Maximum Likelihood (ML) or Bayesian frameworks. The models typically describe the probability of substitution events along a branch of the phylogenetic tree, whereby an amino acid (or nucleotide) is replaced by another. Parameters of these models include rates of substitution, the topology of the phylogeny, and its branch lengths— representing the expected number of substitutions per site occuring along that branch. In the ML framework, parameter inference is achieved by heuristics that attempt to maximize the likelihood. In the Bayesian framework, it is often achieved by Markov Chain Monte Carlo algorithms that sample the posterior distribution. Both approaches are computationally expensive for two reasons. First, they need to explore the space of tree topologies, which grows super-exponentially in the number of leaves [5]. Second, this exploration involves numerous computations of the likelihood, each obtained with a costly sum-product algorithm (Felsenstein’s pruning algorithm [6]). This computational cost has kept researchers from using more realistic models of sequence evolution, which would for instance take into account interactions between sites of a protein (as in e.g., [7]). Such simplifications are well-known to be problematic, as several reconstruction artifacts directly associated to model violations were discovered early in the history of model-based phylogenetic reconstruction [8–10]. Much faster methods exist, but they are generally less accurate [11]. In particular, distance methods (e.g., Neighbor Joining (NJ) [12], BioNJ [13], FastME [14]) build a hierarchical clustering of sequences based on some estimate of their evolutionary pairwise distances, *i.e.*, the sum of the branch lengths along the path between pairs of sequences on the true unobserved phylogenetic tree. NJ is guaranteed to reconstruct the true tree topology if applied to the true distances [15], making the problem of estimating the tree and the set of distances equivalent. In practice, distances are typically estimated under the same probabilistic models as ML and Bayesian methods but considering each pair separately—whereas the latter consider all sequences at once—which greatly simplifies computations but discards part of the global information contained in the full set of homologous sequences.

Here we present Phyloformer, a phylogenetic inference method exploiting all sequences at once with the speed of distance methods. Importantly, Phyloformer can handle complex models of sequence evolution for which likelihood computations would not be feasible. We build on recent advances in deep learning for multiple sequence alignments [MSAs, 16] and in the likelihood-free inference paradigm (Fig. 1). Sometimes referred to as simulation-based inference [17], this paradigm exploits the fact that simulating data under probabilistic models of sequence evolution is computationally affordable, even in cases where computing likelihoods under these models is expensive. Through simulation we sample a large number of phylogenetic trees and MSAs evolved along these trees, given a probabilistic model under which we want to perform phylogenetic inference. We then learn a function that takes an MSA as input and outputs the evolutionary distances between all pairs of sequences on the tree. This function provides a point inference of the *full set of pairwise distances* under the chosen probabilistic model, conditional to the observed MSA. Learning the function is computationally intensive, but once done, Phyloformer can be used in combination with a distance method to reconstruct a tree from an MSA very rapidly, regardless of the complexity of the model of sequence evolution. We show that under the common LG+GC model [18], Phyloformer leads to phylogenies as accurate as state of the art ML methods but runs two orders of magnitude faster. Under more realistic models, *e.g.* accounting for pairwise dependencies between sites, Phyloformer provides more accurate estimates than all other inference methods.

**Fig. 1:**
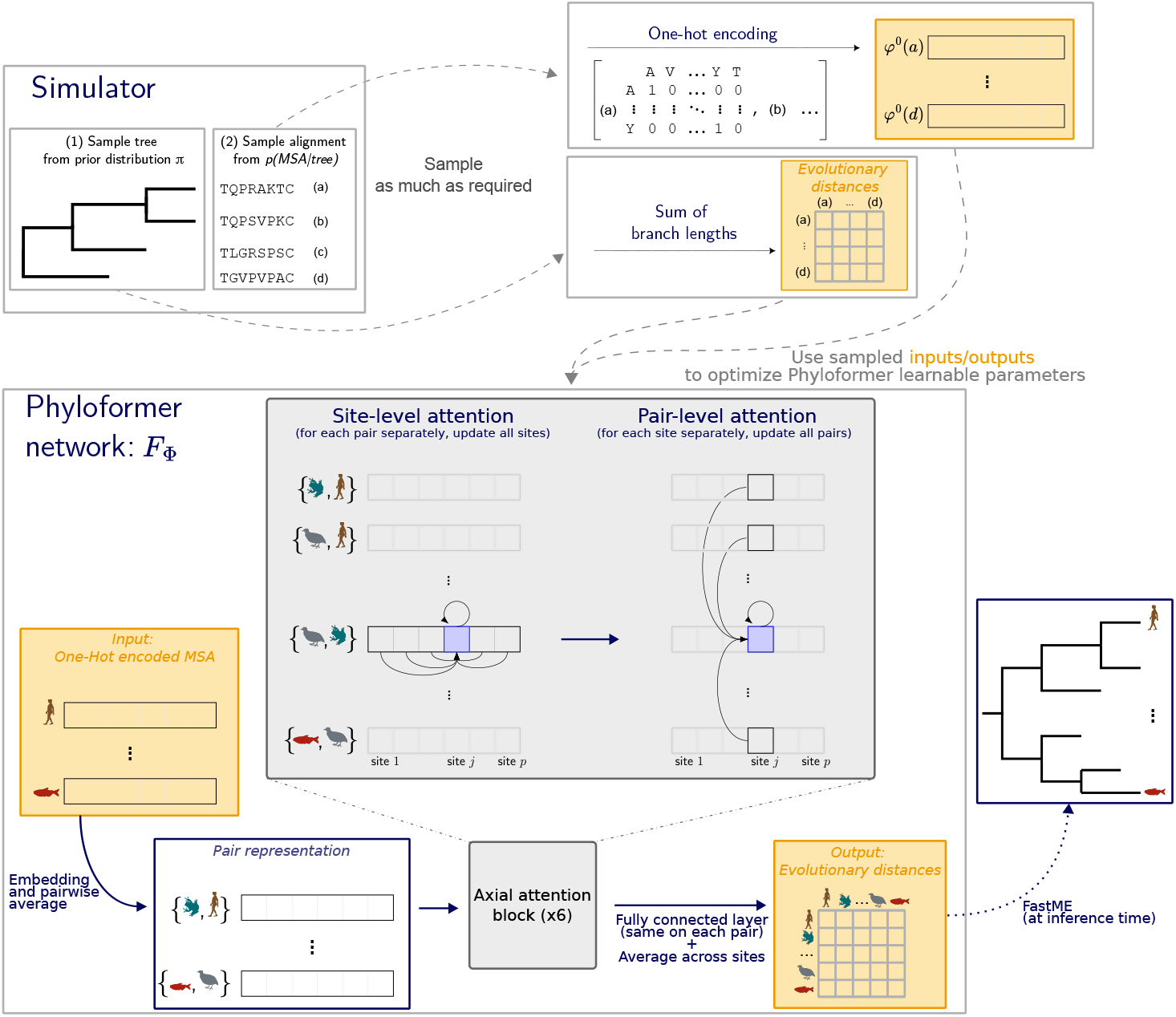
Learning a function that reconstructs a phylogenetic tree from an MSA. We simulate phylogenetic trees and evolve MSAs along these trees under a given probabilistic model (Simulator panel). Once encoded, we use the examples of MSAs and corresponding trees to optimize the prediction function, described in the Phyloformer network panel. Each square denotes a vector of dimension *d* representing one site in one sequence or pair in the MSA, where the value of *d* can be different at each step. Phyloformer starts (bottom left) from a one-hot encoded MSA, and builds a representation for the pairs. These pairs then go through several axial attention blocks which iteratively build a new representation for each pair that accounts for the entire MSA, by successively sharing information across sites within each pair and across pairs within each site (See The Phyloformer neural network). The sharing mechanism relies on self-attention (central panel). We finally use a fully connected network on each site of the resulting representation and average across sites to predict the evolutionary distance between each pair (bottom right). At training time, we compare these distances against real one to optimize the network parameters Φ. At inference time, we feed them to FastME to reconstruct a phylogeny.

**Related work** [19] offer a recent review on deep learning for phylogenetics. [20, 21] proposed likelihood-free methods for phylogenetic inference, by casting the problem as a classification across possible topologies. Given the super-exponential growth of the number of possible unrooted tree topologies in the number of sequences, they restricted themselves to trees with four leaves (quartet trees), that could then be combined to obtain larger trees [22]. Both methods relied on convolutional neural networks and were therefore sensitive to the order of the sequences in the alignment and restricted to a fixed sequence length—smaller sequences being accommodated with padding. More recently, while still only considering quartet trees, [23] proposed a network that was independent of sequence order, and reported accuracies similar to [21] using fewer training samples. [24] showed that the accuracy of the network introduced in [21] was lower than that of ML or distance methods when evaluated on difficult problems involving long branches and shorter sequences (200 sites), for both quartet trees and trees with 20 leaves. [25] proposed a generative adversarial network for phylogenetic inference. While also likelihood-free, this approach required a new training for each inference, and did not scale beyond fifteen species. [26] introduced a distance-based learning method for the related problem of adding new tips into an existing tree. Our work is also related to the recent corpus of methods predicting contact between pairs of residues from MSAs, a crucial step in protein structure prediction [16, 27]. These methods infer distances between sites (columns in the MSA) whereas we infer distances between sequences (rows in the MSA). Our network is trained end-to-end to predict distances, whereas the [16] network is pre-trained on a masked language modeling task to learn a data representation that is then used as input for residue contact prediction learning.

## 2 Results

### 2.1 Likelihood-free phylogenetic inference with Phyloformer

Phyloformer is a learnable function for reconstructing a phylogenetic tree from an MSA representing a set of homologous sequences (Fig. 1). It produces an estimate, under a chosen probabilistic model, of the distances between all pairs of sequences, which is then fed to a fast clustering method to infer a phylogenetic tree. The key feature of Phyloformer is its ability to produce pairwise distance estimates that account for all sequences in the alignment—providing more accuracy than the fast approaches that consider each pair of sequences independently—without computing likelihoods— leading to much faster inference than full ML or Bayesian approaches.

For a given model of sequence evolution *p*(MSA*|τ, θ*) describing how an observed MSA evolves conditionally to a phylogeny *τ* and evolutionary parameters *θ*—substitution rates, equilibrium frequencies—and priors *π*(*θ*) and *π*(*τ*), we generate a large number of samples *{*(MSA*, τ, θ*)*}* under the unnormalized posterior *p*(MSA*, τ, θ*) = *p*(MSA*|τ, θ*)*π*(*θ*)*π*(*τ*) (Fig. 1, Simulator panel). We then use these samples to build a function estimating the tree *τ*, by optimizing a parameterized function *F*_Φ_(MSA) that takes the MSA as input and outputs an estimate of *τ*. More precisely we output point estimates of the distances between pairs of aligned sequences in *τ*, and minimize the average absolute error between these point estimates and the real distances, which amounts to estimating the median of the posterior distribution *p*(MSA*|τ, θ*), see Supplementary Methods 1.4. Assuming that the family of functions described by *F*_Φ_ is expressive enough and that enough samples are used, this approach offers posterior inference under the model (*π, p*), effectively replacing likelihood evaluations by samplings of *p*(MSA*|τ, θ*).

Our *F*_Φ_ relies on self-attention—a mechanism popularized by the Transformer architecture [28]—to build a vector representation for each pair of sequences that contains all the information from the MSA required to determine the corresponding distance. During each self-attention block, the representation of each pair is updated using information extracted from all others. The learnable weights of the block determine how much each pair weighs in the update of any particular pair, as well as what information it contributes. More precisely, we maintain a separate representation for each position within each pair, and alternate between a separate update for each site—whereby information is shared among pairs as we just described—and a similar separate update for each pair whereby information flows among the sites [16, 29]. Following the attention blocks, we use a fully connected neural network on the enriched representation of each pair of sequences to predict the corresponding distance on the phylogenetic tree. The initial representation of each pair is an average of the one-hot encodings of its sequences, that is blind to the rest of the MSA. Because the learnable weights are chosen to make the predicted distances as close as possible to the real ones, we expect them to adaptively extract an MSA-aware representation for each pair, that captures the relevant information from all sequences.

### 2.2 Under a standard model of evolution, Phyloformer is as accurate and much faster than ML

We first assessed the performances of Phyloformer on data generated under the LG+GC model of sequence evolution which combines the LG matrix of amino-acid substitution [18] with rate heterogeneity across sites [30]. The LG model is widely used, implemented in many phylogenetic tools [31–34] and amenable to likelihood computation, making it a good model to compare against state of the art ML inference methods. Following [35], we sampled trees under a birth-death process, subsequently rescaling the branches to simulate variations of the rate of sequence evolution. We chose simulation parameters to match empirical data in the HOGENOM [36] and RaxMLGrove [37] databases (see Online methods). We then evolved MSAs of 50 sequences and 500 sites under LG+GC along these trees, and used the resulting data to train Phyloformer. We compared Phyloformer (PF) followed by FastME to reconstruct the tree from estimated distances against two ML methods, IQTree and FastTree, and one distance method, FastME using LG pairwise distances. Fig. 2a shows the average Kuhner-Felsenstein (KF) distance [38] between the true and reconstructed phylogeny for each of these methods over 500 samples from the same model for increasing numbers of leaves. The KF distance is widely used to compare phylogenies and captures both topological and branch length reconstruction errors. Phyloformer achieved a performance similar to ML methods. It is noteworthy that this performance was stable across numbers of leaves, even though our network was trained on 50-leave phylogenies only. The performance was also stable when doing inference over a range of sequence lengths, even though Phyloformer was trained only on alignments with 500 positions (Supplementary Figs. 13 and 14). The distance method was much less accurate. Interestingly, the high accuracy of Phyloformer was achieved with the lowest runtime among all benchmarked methods (Fig. 3). In particular, it was up to 135 times faster than the ML method IQTree, for a similar accuracy. FastTree—a faster and supposedly less accurate heuristic for ML—also had similar accuracy on this dataset, but remained one order of magnitude slower than Phyloformer. Phyloformer was even twice as fast as FastME combined with LG distances. As Phyloformer itself runs FastME to reconstruct a tree from its distance estimates, this difference indicates that inferring distances that exploit the full MSA with a trained Phyloformer on a GPU is actually faster than computing the ML distances independently for each pair. Conversely, Phyloformer was the most memory intensive method, using up to 7.4GB of GPU RAM (Supplementary Fig. 15), although this can be halved in some cases by using automatic mixed precision.

**Fig. 2:**
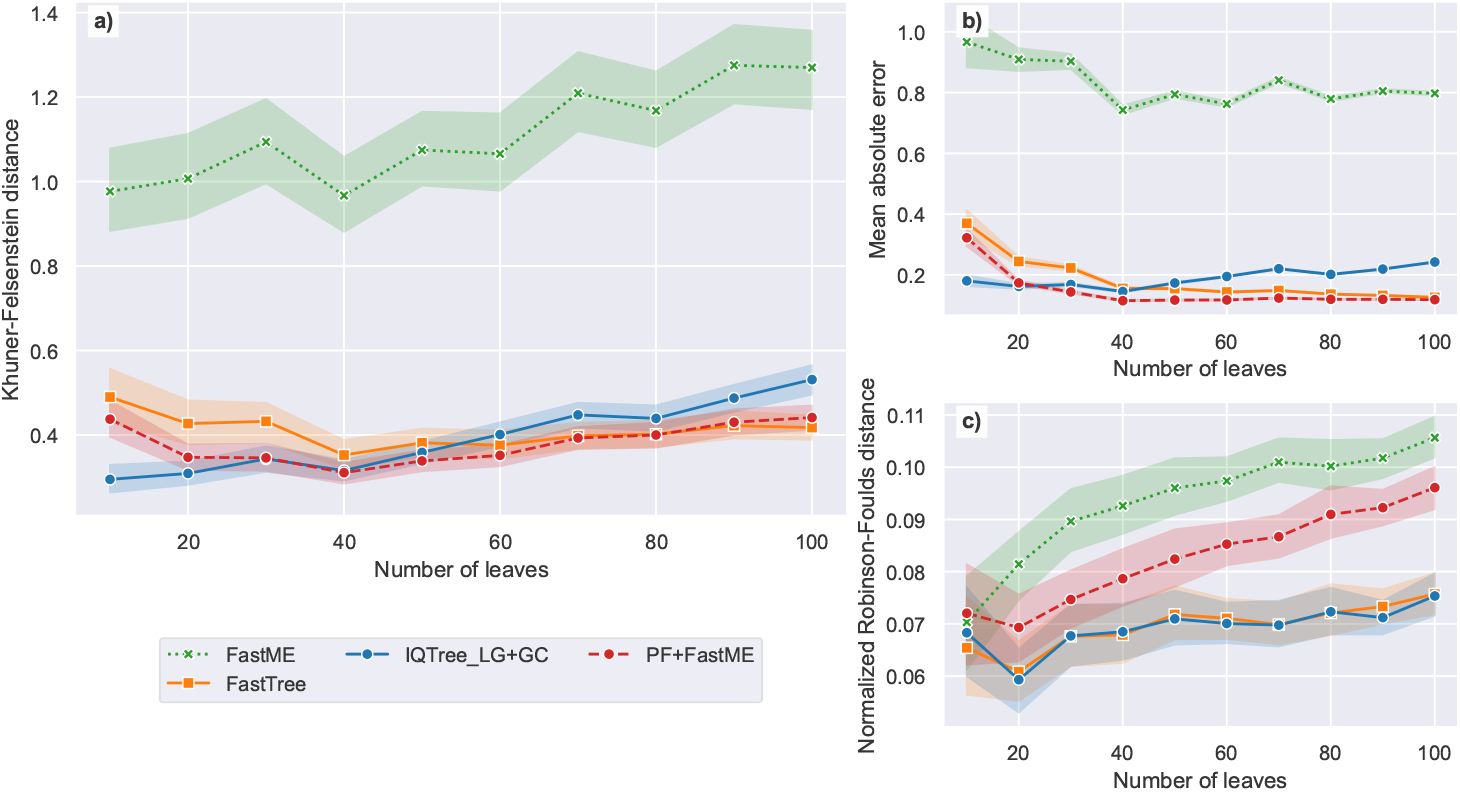
Performance measures for different tree reconstruction method. **a)** Kuhner-Felsenstein (KF) distance, which takes into account both topology and branch lengths of the compared trees; **b)** mean absolute error (MAE) on pairwise distances, which ignores topology; **c)** normalized Robinson-Foulds (RF) distance, which only takes into account tree topology. The alignments for which trees are inferred, were simulated under the LG+GC sequence model and are all 500 amino acids long. For each measure, we show 95% confidence intervals estimated with 1000 bootstrap samples.

**Fig. 3:**
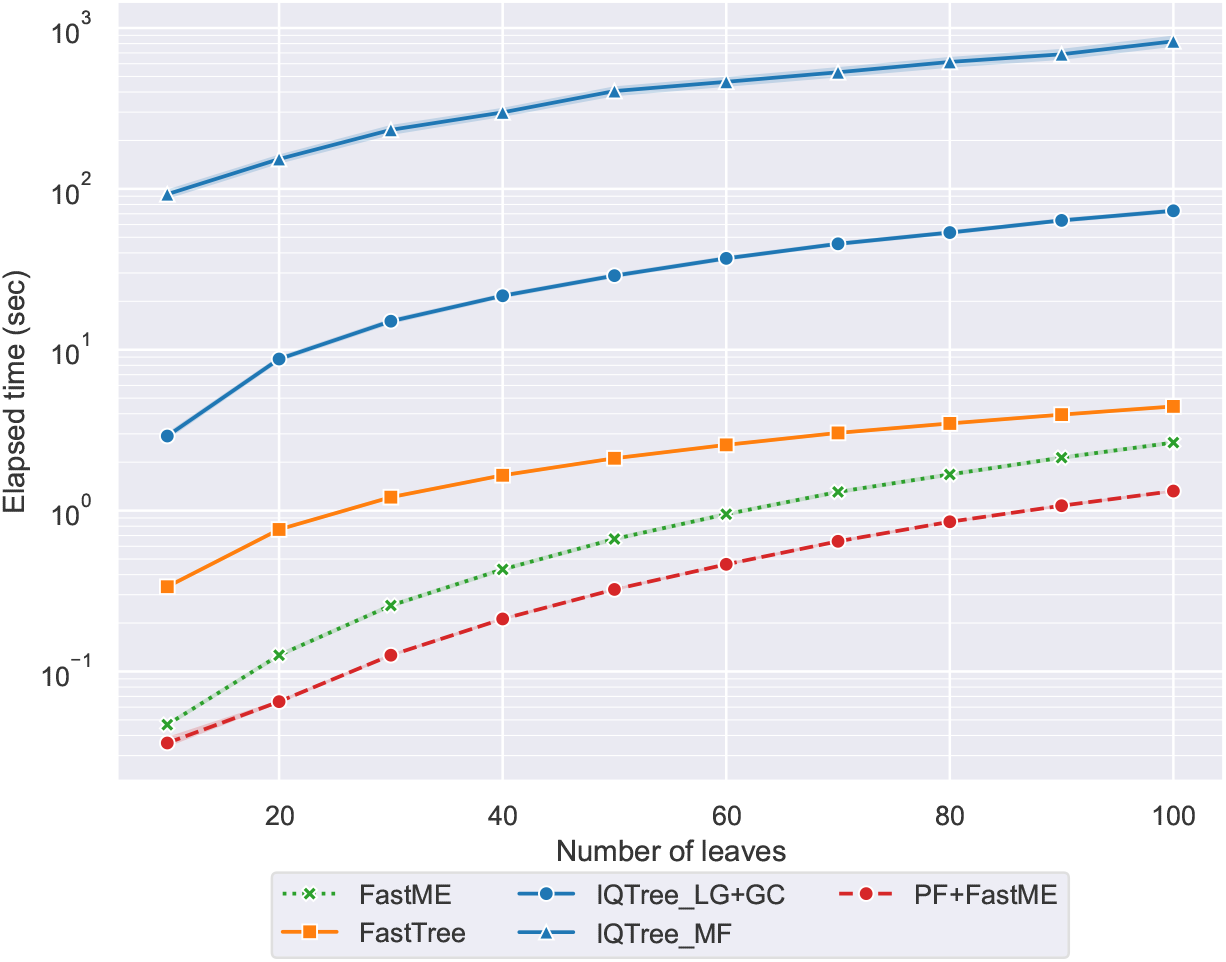
Execution time for different tree reconstruction methods on the LG+GC test set with alignments of length 500. For IQTree ModelFinder (MF) times were measured on the Cherry testing set (see Section 2.3). For all methods except Phyloformer, total wall time was measured. For Phyloformer, the elapsed time is the sum of the time it takes to infer the distances and the time FastME takes to infer the tree from these distances. It is important to note that the distance prediction time does not include the time it takes to load the Phyloformer weights to the GPU as we did that once before inferring distances for all the testing alignments.

Fig. 2b&c stratify the reconstruction error in terms of their topology (panel c, using the normalized Robinson-Foulds (RF) metric [39]) and pairwise distances (panel b, using the Mean Absolute Error (MAE) between true and estimated distances). Regardless of the criterion, Phyloformer dominated FastME by being both faster and more accurate. On the other hand, Phyloformer reconstructed topologies that were less accurate than ML methods, and increasingly so for larger numbers of leaves, but estimated distances as or more accurately. A possible explanation for this discrepancy is that since we control the tree diameter in our simulation, larger trees have shorter branches on average. As branch lengths decrease, the number of mis-predicted branches increases leading to larger topological errors (see Supplementary Results 2.3 for an in-depth explanation).

Finally, we investigated the ability of Phyloformer to handle gaps contained in empirical MSAs because of insertion-deletion (indel) events that have occurred during sequence evolution. Standard models of sequence evolution consider gaps as wildcard ‘X’ characters, and thus cannot benefit from the information they provide. Models that account for insertion-deletion processes are more complicated to implement and more costly to run [40], but can easily be included using our paradigm. We fine-tuned the Phyloformer network previously trained on ungapped LG+GC data on a smaller dataset that includes indels, inserted through a model of insertion/deletion events in Alisim [41], choosing parameters as in [42]. Fig. 4 shows that the accuracy of all methods dropped on alignments that include gaps compared to alignments that do not (Fig. 2), probably because gaps remove information from the alignments. However the difference between Phyloformer and ML methods shrinked, with Phyloformer outperforming ML methods according to the RF metric for 10 to 30-leaf trees. This is likely due to Phyloformer’s ability to extract information from gaps, which are encoded as a separate character and not as a wildcard character.

**Fig. 4:**
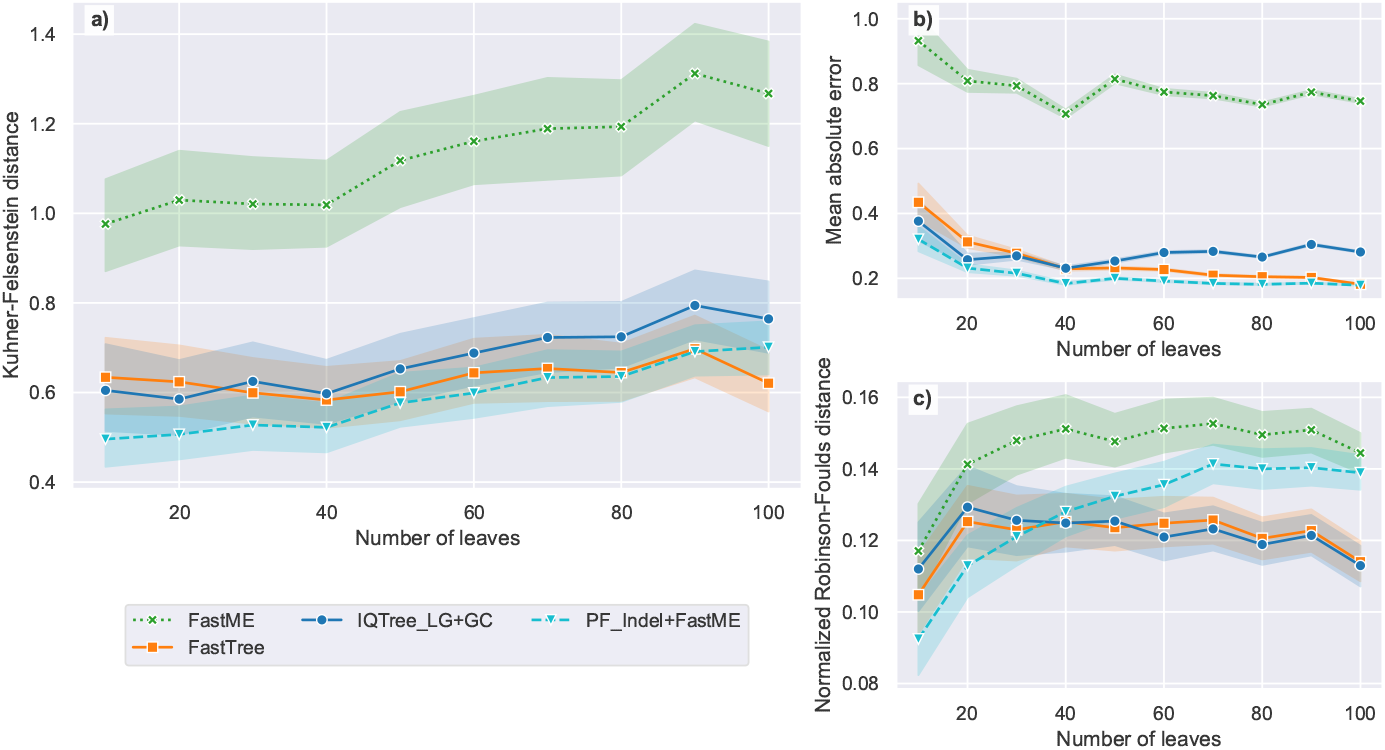
Tree comparison metrics for different tree reconstruction methods on the LG+GC+indels test set (alignment length=500). Legend as in Fig. 2, with Phyloformer finetuned on alignments with gaps named PF_Indel_+FastME and in cyan.

### 2.3 Under more realistic models, Phyloformer outperforms all other inference methods

Because ML and Bayesian inference approaches must compute the likelihood, in practice they can only be used under simple models such as LG+GC for which these likelihood calculations are affordable. Phyloformer on the other hand can reconstruct phylogenies under arbitrarily complex models of sequence evolution, as long as we can efficiently sample training data from these models. We now illustrate this feature by considering inference tasks under two substitution models that relax common simplifying assumptions: independence between sites, and the homogeneity of selective constraints across sites. The first model we used (Cherry, Supplementary Methods 1.2) is derived from a model of sequence evolution that includes pairwise amino-acid interactions [43]. ML inference under such a model would be very costly for two reasons: the substitution matrix has size 400 *×* 400, and would need to be applied to pairs of interacting sites, which would need to be identified with additional computations. The second model (SelReg, Supplementary Methods 1.2) draws different selective regimes for each site of the alignment: a site can evolve under neutral evolution, negative selection, or persistent positive selection. ML inference under such a model is achievable with a mixture model [*e.g.*, 44], but costly, because the SelReg mixture includes 263 distinct amino acid profiles, plus a profile for neutral evolution, and a different matrix for positively selected sites. We fine-tuned the Phyloformer network previously trained under the LG+GC model on alignments sampled under the Cherry or the SelReg model. We compared its performances against the same methods as before, but allowing IQTree to search for the best evolution model available (with the Model Finder option). Fig. 5 shows that under both the Cherry and SelReg models all methods performed worse than under LG+GC, presumably because both models decrease the information provided by a given number of sites, by including pairwise correlations (Cherry), or positively selected sites that are likely to saturate (SelReg). However, Phyloformer outperformed all other methods by a substantial margin, with KF distances around 1 whereas others range between 2 and up to 10 for IQTree under SelReg. Of note, the Model Finder option was costly, further increasing the computational edge of Phyloformer (Fig. 3). Not using this option markedly decreased the accuracy of IQTree on the Cherry alignments (Supplementary Figs. 11 and 12). As we observed under LG+GC, Phyloformer was better at estimating distances than topologies (Fig. 5), with the latter becoming more challenging for larger numbers of leaves. Nonetheless, the RF distances of Phyloformer remained lower than for other methods, except for SelReg on trees with more than 40 leaves where it was only outperformed by IQTree—which in turn had the worst distance estimates among all benchmarked methods.

**Fig. 5:**
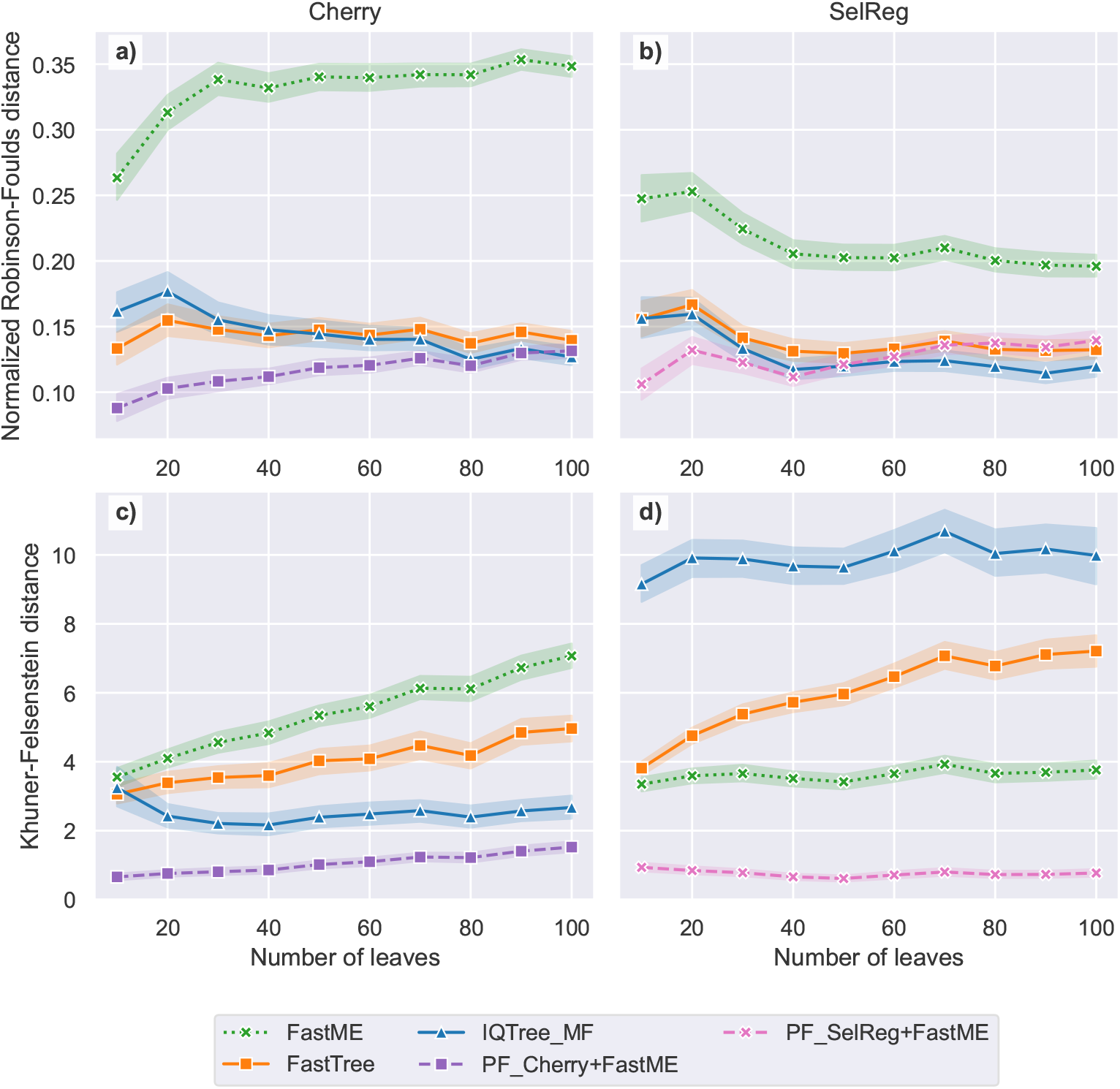
Normalized Robinson-Foulds distance (above) and Kuhner-Felsenstein distance (below) for different tree reconstruction methods on the Cherry (left) and SelReg (right) test sets (alignment length=500).

### 2.4 Phyloformer performs on par with ML methods on empirical data

We compared the performance of Phyloformer and other methods on 346 orthologous gene alignments from 36 Cyanobacteria [45], reasoning that good reconstruction methods should more often infer trees that match the tree obtained on the concatenated gene alignments. We compared the LG+GC-with-indel version of Phyloformer to the same three methods assessed in section 2.2. Fig. 6a shows that Phyloformer performed as well as the other standard methods on empirical data, and did so faster.

**Fig. 6:**
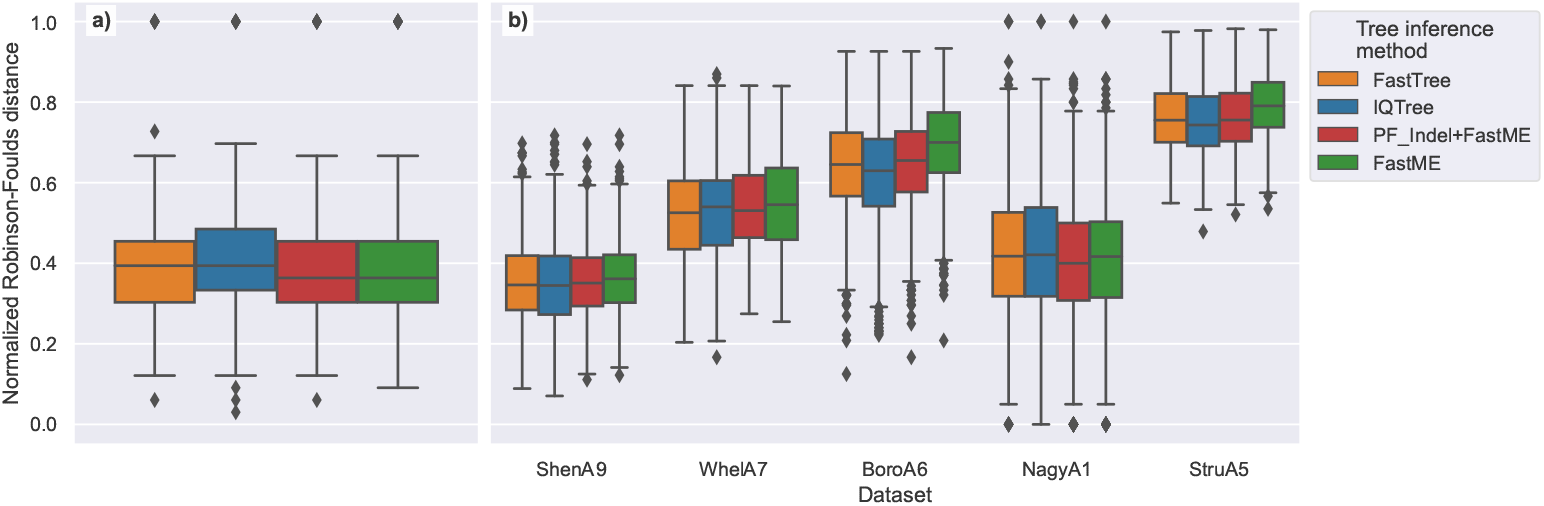
Comparison of topology reconstruction accuracy between Phyloformer and other methods on empirical data. In both panels, we show the normalized Robinson-Foulds distance between reconstructed gene trees and the corresponding concatenate tree. In **a)** inferred gene trees on alignments from [45] using the same pipeline as in section 2.2 and with the gap-aware version of Phyloformer shown in Fig. 4. In **b)** gene-alignments, species trees and some gene trees were obtained from [46]. We inferred gene-trees using the gap-aware version of Phyloformer and FastME as in panel **a)**. The IQTree predictions were made in [46] under the evolutionary model found by IQTree-ModelFinder, then 10 predictions were done and only the one with the best likelihood was kept. The datasets shown here, have *≥* 80% of alignments detected as LG by IQTree.

We conducted a similar analysis on gene-trees over many different species and orders obtained from [46]. In this study the authors collected a large number of sequence datasets and inferred gene-trees using IQTree and FastTree under the evolutionary model found by IQTree’s ModelFinder for each alignment. For IQTree they inferred 10 trees and only kept the one with the best likelihood. The authors also reconstructed species trees from concatenated alignments for each dataset. We reconstructed trees on the gene families where at least 80% of alignments were classified as LG by IQTree using the LG+GC-with-indel version of Phyloformer with FastME. We then compared our gene trees as well as the ones from [46] to the concatenate trees. Here again, Fig. 6b shows that in most cases Phyloformer performed as well as the best of 10 trees estimated with ML methods. Here the computational speed of Phyloformer shines as we were able to infer about 12, 000 trees in under two hours with one GPU. In [46], the authors measured execution times of only 10% of tree inference tasks, for which the total runtimes of IQTree and FastTree were approximately 10.5 days and 4 hours respectively. On the same subset of trees, we measured the total runtime of Phyloformer+FastME and standalone FastME at approximately 11.5 and 15 minutes respectively. Furthermore, Phyloformer consistently produced trees with a higher likelihood than FastME trees though still lower than pure ML methods (Supplementary Fig. 7).

## 3 Discussion

Drawing on recent breakthroughs in likelihood-free inference and geometric deep learning, we have demonstrated that Phyloformer achieves rapid and precise phylogenetic inference. The likelihood-free paradigm only requires samples from the probabilistic model of sequence evolution, which allows inference under much more complex models than ML or Bayesian inference. Furthermore we exploited an amortized form of this paradigm, requiring a single training of a neural network that takes an MSA as input and outputs evolutionary distances between pairs of sequences—as opposed to approaches like ABC [47] that require a new sampling step at each inference. We based our neural network on axial self-attention, an expressive mechanism that accounts for the symmetries of the MSA and seamlessly handles arbitrary numbers of sequences of any length.

Phyloformer was faster and as accurate as ML inference methods on data sampled under the standard LG+GC model. Computing likelihoods under LG+GC is expensive but possible, making ML inference the gold standard: reaching the same accuracy faster was the best outcome one could hope for. On the other hand, computing likelihoods under more complex models accounting for local dependencies (Cherry) or heterogeneous selective pressures (SelReg) is too costly, forcing ML methods to work under misspecified models whereas Phyloformer can still perform inference under the correct model, without any effect on its speed. As a result, Phyloformer yields the most accurate inference by a substantial margin while retaining its computational edge.

More generally, we stress that likelihood-free inference using neural networks has a model-based nature identical to ML or Bayesian methods. It formally estimates the posterior distribution defined by the prior and probabilistic model used to simulate training data, accessing this model through sampling instead of likelihood evaluations. As such, it is not immune to model misspecification: we observed for example that Phyloformer trained on LG+GC underperformed on data simulated under Cherry or SelReg and vice-versa (Supplementary Figs. 11, 12 and 16). Rather than replacing model choice, we believe that the crucial contribution of a likelihood-free method like Phyloformer is to offer a way to work under more realistic models of sequence evolution that were so far not amenable to inference.

It is noteworthy that the inference speed that we report for Phyloformer was recorded on a GPU, a less widespread hardware than the CPU used for other methods, which may limit its interest for analysing a single gene alignment under models amenable to ML. However, we expect Phyloformer to have a significant impact in experiments where many reconstructions are necessary, *e.g.* for bootstrapping, reconstructing several gene trees from whole genomes or transcriptomes, or where more complex models are warranted. Another current limitation of Phyloformer is its scalability. The current bottleneck is on its memory usage, mostly driven by applying self-attention to pairs of sequences. A better scaling version could be obtained by working at the sequence level— attempts to do so have so far led to lower accuracies. An important extension of Phyloformer will be to train with a topological loss function, *e.g.* directly minimizing the RF metric rather than a distance metric. Such a version would address the gap that we observed between accuracies in distance and topological reconstruction, and could also lead to a more scalable method by working around the need for all pairwise distances—of quadratic size in the number of sequences whereas the tree itself has linear numbers of nodes and edges. We also believe that extending Phyloformer to unaligned sequences will be of interest, both because multiple alignments are computationally intensive, and because they are error-prone. This could be addressed by including the alignment step in the network [48, 49]. Alternatively, one could forego alignment altogether, *e.g.* by producing a length-independent representation early in the neural network.

We expect that Phyloformer will have its largest impact on phylogenetic inference after versions are trained on a collection of more realistic models of sequence evolution which could include nucleotides, variations along the sequence or between branches and position-specific dependencies among sites [7, 50, 51]. Our self-attention network could exploit these latter dependencies via the addition of positional encodings—a standard approach in the transformers literature. Beyond phylogenetic reconstruction, our network can be trained to infer other parameters of the simulation model. This would provide an efficient and flexible way to study phylodynamics, phylogeography, and selective pressures operating on the sequences, for instance.

## 4 Online methods

### The Phyloformer neural network

Phyloformer is a parameterized function *F*_Φ_ that takes as input an MSA of *n* sequences of length *L* and outputs an estimate of the *N* = (*^n^*_2_) distances between all pairs of sequences. Φ denotes the set of learnable parameters of *F*_Φ_. We then input these distances to FastME [14] to obtain a phylogenetic tree (Fig. 1).

The Phyloformer network starts with a one-hot encoding of the aligned sequences: every sequence *x* is represented as a matrix *φ*^(0)^(*x*) *∈ {*0, 1*}*^22×^*^L^* in which column *j* contains a single non-zero element 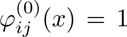, whose coordinate *i ∈ {*1*,...,* 22*}* denotes the amino acid or gap present in sequence *x* at position *j*. It then represents each pair (*x, x*^′^) of sequences in the MSA by the average of their individual representations *i.e.*, with a slight abuse of notation,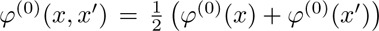. Of note, *φ*^(0)^(*x, x*^′^) does not depend on the order of sequences *x* and *x*^′^. At this stage, the network represents each site within each pair independently of all others, encoding information such as “at site 4, sequences *x* and *x*^′^ contain a Leucine and an Isoleucine”. The whole purpose of *F_ϕ_* is to account for relevant information about the evolutionary distance between *x* and *x*^′^ contained in other sequences from the alignment. To extract this information, *F_ϕ_* uses *r* = 6 self-attention layers [28] that iteratively build updated *φ*^(^*^l^*^)^(*x, x*^′^) *∈ d × L* representations of each pair using all others in the MSA. More precisely, we use axial attention [16, Fig. 1, central panel] and successively update each pair (resp. site) separately by sharing information across sites (resp. pairs). Along each axis, we rely on a modified linear attention [52, see Scalable self-attention], with *h* = 4 attention heads and embeddings of dimension 64 for the value matrix and only 1 for the query and key matrices. The *r* axial attention blocks of Phyloformer output for every pair of sequences a tensor *φ*^(^*^r^*^)^(*x, x*^′^) *∈* R*^d^*^×^*^L^* informed by all other pairs in the same MSA. We convert this representation into a single estimate of the evolutionary distance between *x* and *x*^′^ by applying an R*^d^ →* R fully connected layer to each site of each pair, followed by an average over the sites. We provide more details on the *F*_Φ_ architecture in Supplementary Section 1.3.

### Accounting for symmetries

It is now well understood that accounting for known symmetries is key to the success of deep learning, as formalized in geometric deep learning [53]. Following this principle, we parameterize the function *F*_Φ_ by a neural network that exploits two symmetries of the estimation task: the estimated evolutionary distances should not depend on the order of the *n* sequences or *L* sites in the MSA. More precisely, we want *F*_Φ_ to be equivariant by permutations of the sequences: if it returns values *d_ab_, d_ac_, d_bc_* when presented with sequences (*a, b, c*), it should return *d_ac_, d_bc_, d_ab_* when given (*c, a, b*) as input. On the other hand, *F*_Φ_ should be invariant to permutations of the sites—any such permutation should lead to the same *F*_Φ_ distances. This last point may seem counterintuitive as the order of residues in a protein matters for its function, and it is known that close residues do not evolve independently. Nonetheless in all our experiments we train—or pre-train—*F*_Φ_ on data generated under the LG+GC model, which is site-independent. The self-attention updates act on the R*^d^* representations of a site within a pair of sequences regardless of their order, yielding the desired equivariances. Enforcing these equivariances would be more difficult if the updates were general functions acting on entire MSAs represented by R*^d^*^×^*^N^*^×^*^L^* tensors. The final average across sites within each pair makes *F*_Φ_ invariant rather than equivariant by permutation of these sites. In addition because none of the operations in *F*_Φ_ depend on the number of sites or pairs, we can use the same *F*_Φ_ seamlessly on MSAs with an arbitrary number of sequences of arbitrary length.

### Scalable self-attention

Naive implementations of self-attention over *M* elements scale quadratically in *M* —in our case, both the number of sites and pairs of sequences. Indeed, softmax attention as introduced by [28] is parameterized by three matrices *Q, K, V ∈* R*^M^*^×^*^d^* for some embedding dimension *d*, respectively called Queries, Keys and Values, and every update for an element *i* computes attention weights (*s_i_,1,...,s_i_,M*) = softmax 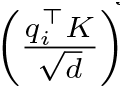. We resorted to the linear attention of [52], who exploited the fact that *s_ij_* = 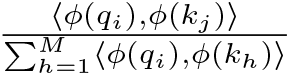 for some non-linear infinite-dimensional mapping *ϕ* : R*^d^ → H* to a Hilbert space *H* [54] and proposed to replace *ϕ* by some other non-linear, finite-dimensional mappings *ϕ̃* : R*^d^ →* R*^t^*. We can then re-write the self-attention updates 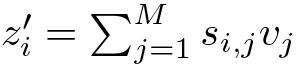 as

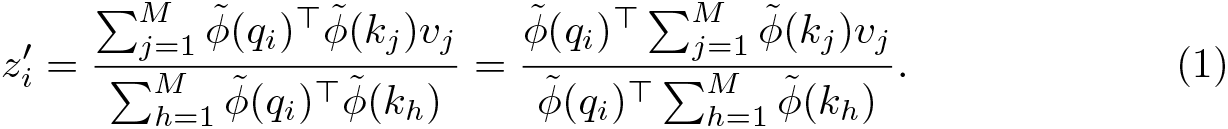

Because we can pre-compute each of the two sums and re-use it for every query, this simple factorization reduces both the number of operations and memory usage from *O*(*M* ^2^ *· L · d*) to *O*(*M · L · d · t*). Following [52] we used an ELU-based mapping [55]

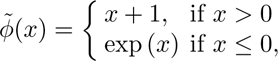

where the operation is applied entrywise, yielding *ϕ̃*(*x*) *∈* R*^d^* vectors for *x ∈* R*^d^*. In our experiments, we used *d* = 64 for the Values matrix, but noticed that using *d* = 1 for Queries and Keys led to slightly lower training-loss values (Supplementary Fig. 20a), while substantially reducing the memory footprint of the self-attention layers (Supplementary Fig. 20b). This observation is consistent with recent research showing that Transformers and other neural networks learn through gradual rank increase [56, 57]. However, applying (1) with queries and keys of dimension 1 leads to identical updates *z_i_*^′^for all elements. To work around this issue, we normalized each update by the average of queries and the sum of keys instead of the usual sum of attention weights, leading to

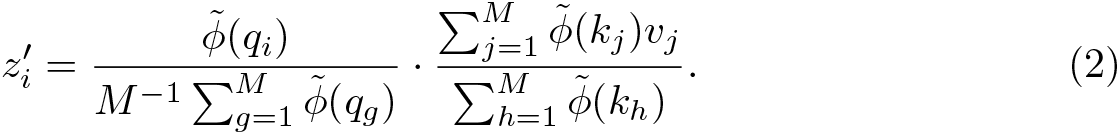

### Training Phyloformer

We trained *F*_Φ_ using 6 NVIDIA A100 80GB GPUs on simulated examples through a loss function (see Metrics) comparing the estimated and true evolutionary distance (Fig. 1). We used the Adam optimizer [58], batches of size 4 and a maximum learning rate of 10^−3^ with 3000 linear warmup steps followed by a linear decrease of 213,270 steps, corresponding to 30 epochs. We also implemented an early stopping criterion that stopped training when the validation loss did not decrease over 5 successive 3000 step intervals.

We first trained an *F^pre^*_Φ_ function that served as a starting point for all the functions used in our experiments, by optimizing Φ with respect to the MAE loss for 20 epochs (*≈* 79 hours) over the 170,616 examples (see Section 4) simulated under LG+GC, saving a model every 3000 steps, and eventually retaining the one with lowest Robinson-Foulds error (see Metrics) over the validation dataset (17016 examples). For the results in Fig. 2, we further optimized the parameters of *F^pre^*_Φ_ for 4 epochs (20 hours) with respect to the MRE loss leading to a slightly improved error over small distances (Supplementary Fig. 17) and on the overall Robinson-Foulds metric (Supplementary Fig. 10). For the results in Figs. 4 and 5 we further optimized the parameters of *F^pre^*_Φ_ for the MAE loss on gapped MSAs and MSAs generated under the Cherry or SelReg substitution models respectively (see Datasets).

### Baselines

*IQTree LG+GC* [59, v2.2.0] reconstructs phylogenies in the Maximum Likelihood framework. It first estimates several parsimony trees along with one reconstructed through a distance method, then optimizes branch lengths and other parameters of the model of sequence evolution, while performing local topological rearrangements (Nearest Neighbor Interchanges, NNIs) to maximize the likelihood. We ran it with the LG model of amino acid substitution [18] combined with a continuous gamma distribution to model rate heterogeneity across site [60]. In our experiments we did 5 rounds of NNIs since we observed that optimizing for more rounds rarely improved the topology of the final tree while substantially adding to the running time. The software was run with iqtree2 -T 1 -m LG+GC -n 5.

*IQTree MF* uses the Model finder (MF) mode of IQTree [61], in which likelihoods of an initial tree are computed for a large set of substitution models and models of rate-heterogeneity accross sites. The best fitting model is selected using BIC. The rest of the tree search is done as above but using the selected model for likelihood estimations. The software was run with iqtree2 -T 1 -n 5.

*FastTree* [62, v2.1.11 SSE3] reconstructs a starting tree using an algorithm inspired from Neighbor-Joining [12] which is subsequently refined with topological rearrangements to optimize the minimum evolution criterion. The tree is then improved using maximum likelihood with NNIs. It was run under the LG+G4 model of sequence evolution. The software was run with fasttree -lg -gamma.

*FastME* [14, v2.1.6.4] computes a distance matrix using Maximum Likelihood, then reconstructs a tree topology using BioNJ [63] and further refines it via topological rearrangements which seek to optimize the Balanced Minimum Evolution score. In virtually all performed experiments we observed that the FastME tree search algorithm led to slightly better performances than the neighbor joining algorithm [12]. We didn’t resort to the --gamma option as in our experiments we observed that this lead to worse performances. Using FastME as our baseline distance method makes the comparison with Phyloformer insightful, as the only difference between the two methods is the distance matrix used as input. The software was run with fastme --nni --spr --protein=LG to reconstruct trees using the inbuilt evolutionary distance estimation and simply with fastme --nni --spr when Phyloformer’s predicted distance matrix was provided.

All methods were run on a single CPU thread (Intel Xeon E5-2660 2.20GHz) except for Phyloformer distance prediction which was run on a single GPU (NVIDIA V100 32GB).

### Datasets

We generated ultrametric phylogenies under a birth-death process. We used 50-leaf trees for training, and 10-leaf to 100-leaf trees for testing. We rescaled branch lengths as in [35] to yield non-ultrametric trees. Finally, we rescaled each tree to resemble trees found in public empirical databases. We used each rescaled phylogeny to simulate one MSA with AliSim [41] for the LG+GC model, or in-house code for Cherry, or Pastek [64] for SelReg. For LG+GC, we sampled the parameter of the gamma distribution to match values estimated on empirical data. We provide more details in Supplementary Methods 1.

### Metrics

We now describe the metrics used throughout this article to compare phylogenies or optimize our network.

Let *d_i_* be the *i*^th^ of *N* true evolutionary distances in a phylogeny, and *d*^^^*_i_* the corresponding estimate output by a given tree inference method. Then the mean absolute error (MAE) and mean relative error (MRE) are defined as

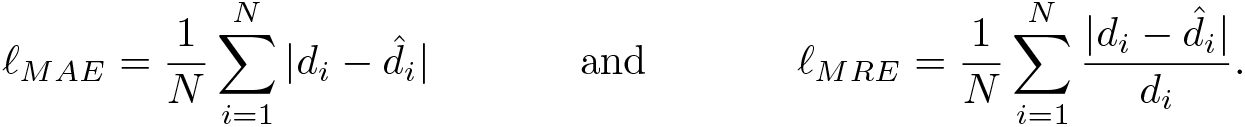

When used to compute the loss during Phyloformer training, *d*^^^*_i_* values correspond to distance estimates directly output by *F*_Φ_. When used as a metric (e.g. in Fig. 2) we use *d*^^^*_i_* values extracted from the reconstructed tree, by summing all branch lengths on the paths between each pair of leaves—even for Phyloformer— in order to fairly compare different methods.

In phylogenetic trees, each branch describes a bipartition of the set of leaves, paired with a weight (*i.e.*, the branch length). Let *A* and *B* be the sets of leaf-bipartitions describing trees *T_A_* and *T_B_*, and *w_e,T_* the weight of a bipartition *e* in tree *T*. Then, the Normalized Robinson-Foulds distances and the Kuhner-Felsenstein distance between *T_A_* and *T_B_* can be written

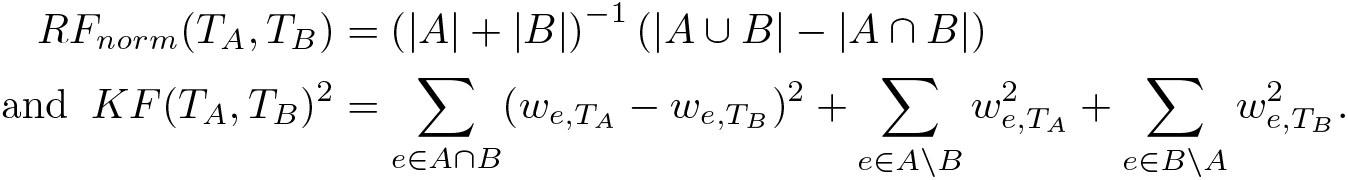

## Code and data availability

The code for Phyloformer, the pretrained models, and all the datasets analyzed in this work can be found at https://github.com/lucanest/Phyloformer.

## Acknowledgements.

The authors thank Dexiong Chen, Flora Jay, Martin Ruffel, Johanna Trost for insightful discussions.

This work was funded by the Agence Nationale de la Recherche (ANR-20-CE45-0017). It was granted access to the HPC/AI resources of IDRIS under the allocation AD011011137R1 made by GENCI. Part of this work was performed using the computing facilities of the CC LBBE/PRABI. The taxon silhouettes in Fig. 1 are modified from public domain images in the PhyloPic database.

