## Supplementary material for "Phyloformer: Fast, accurate and versatile phylogenetic reconstruction with deep neural networks"

**3 Supplementary Figures** **10**

#### 1 Supplementary Methods

Here we provide a detailed description of our protocol to sample trees and multiple sequence alignments along these trees.

##### 1.1 Simulating phylogenies

We chose our simulation parameters to generate trees similar to empirical tree distributions.

##### 1.1.1 Empirical tree distributions

We collected $63,245$ phylogenies from the HOGENOM and $4,893$ phylogenies from the RaxMLGrove databases. In order to have the same order of magnitude across datasets we multiplied by 10 the frequency of the latter in the distribution. We furthermore kept all the trees with diameters in the range $(0.02, 15)$, discarding $9,055$ trees out of $112,175$. The trees sampled from the HOGENOM database correspond to the rooted "nocore" trees from the phyla Delta-Epsilon ($7,687$ trees), Alpha-proteobacteria ($14,125$ trees), Beta-proteobacteria ($11,961$ trees), Tenericutes ($1,634$ trees), Archaea ($7,658$ trees), Spirochaetes ($3,600$ trees), Cyanobacteria ($4,898$ trees) and Bacteroidetes-Chlorobi ($11,682$ trees). The trees sampled from RaxMLGrove correspond to all the trees reconstructed from MSAs of amino acid sequences.

##### 1.1.2 Simulating trees

We trained all neural networks presented in our work on trees/alignments with 50 leaves/sequences, and considered smaller and larger numbers for testing. We followed the same procedure for all trees. Following [1] we simulated each tree using a birth-death process which returns an ultrametric tree using dendropy's `treesim.birth_death_tree` method [2, v4.6.1], then obtained a non-ultrametric tree by rescaling the branches of the tree. To do so we simulated changes of the rate of evolution by running a process that generates small and big rate changes from the root of the tree to the leaves. Then we rescaled all the trees, irrespective of the number of leaves, to match the distribution of diameters (maximum pairwise distance in the tree) observed in empirical data (subsection 1.1.1). For each tree we sampled a diameter from this distribution, and added gaussian noise to provide variation ($\mu = d, \sigma = d/10$, with $d$ the sampled diameter).

Given that all trees share the same diameter distribution irrespective of their number of leaves, trees with more leaves will have a larger number of short branches. This leads to a distribution shift between training and testing data (see Supplementary Fig. 9a). Moreover very short distances also lead to duplicate sequences during the alignment simulation step. To mitigate this and avoid too many alignments with duplicate sequences, we further rescaled terminal branches, resampling their length $l$ from a normal distribution, centered around the minimum allowed valued $\mu = 0.001$, and having $\sigma = 0.005$, until we reached $l \geq \mu$. Supp. Fig. 1 illustrates the whole tree simulation pipeline.

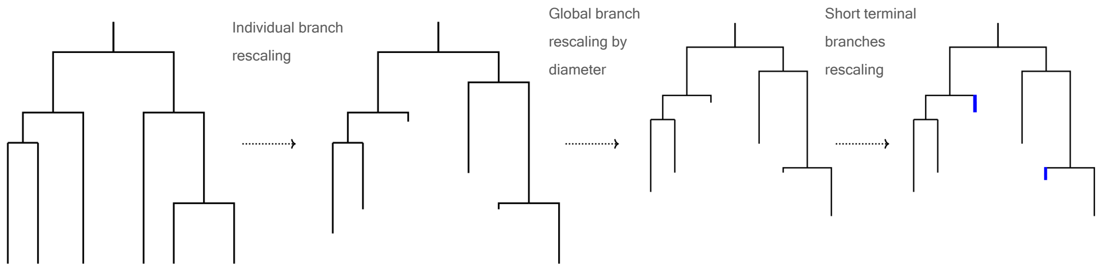

Figure 1: Tree simulation pipeline: we simulate an ultrametric tree with a birth-death process, rescaling the branches individually to simulate changes in the rates of evolution, then rescaling all the branches to match a diameter sampled from empirical data, and finally rescaling terminal branches which are too short to make them longer than the minimum allowed value.

##### 1.1.3 Comparison of simulated and empirical trees

To make sure the simulations were realistic enough we computed several statistics of the training trees: mean distance between leaves, standard deviation of distances, maximum distance, mean branch length, standard deviation of branch lengths, maximum branch length, mean root to leaf distances, standard deviation of root to leaf distances. We compared these statistics to those of the trees extracted from the empirical databases. The distributions of the statistics were markedly different between the two databases (Supplementary Fig. 2) and we chose simulation parameters to overlap with both.

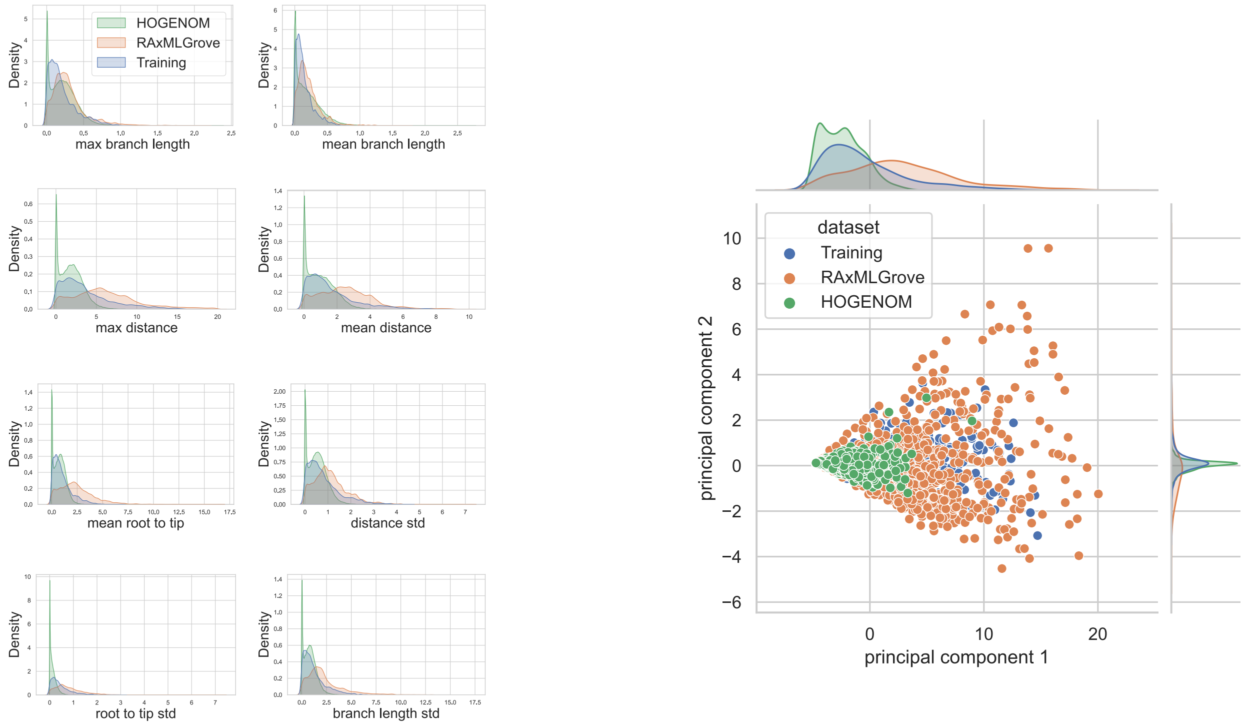

Figure 2: The distribution of trees used to train and test our networks is similar to the empirical tree distributions. Left: Distributions of the considered tree statistics for the empirical tree datasets and the training simulations. The blue curves correspond to simulated trees, the orange curves to RAxMLGrove trees, and the green curves to HOGENOM trees. Right: Two component PCA based on all 8 statistics for the three datasets (1000 points per dataset shown).

##### 1.2 Simulating multiple sequence alignments

In the following we describe how we simulated alignments along the generated phylogenies. Given that the whole tree+alignment simulation pipeline still leads to some alignments having duplicate sequences, we simulate a larger number of data points than we need and discard alignments with duplicated sequences and the corresponding trees.

**LG+GC** We generated alignments under the LG+GC model of evolution along the trees with the Alisim software implemented in IQTree2. We sampled the $\alpha$ parameters of the continuous gamma distribution from those inferred by IQTree on $12,408$ alignments from the HOGENOM core database, and added gaussian noise to ensure variation ($\mu = \alpha, \sigma = \alpha/10$, with $\alpha$ the sampled value). A threshold of $0.05$ was set for the minimum allowed value.

**LG+GC+Indels** We simulated alignments as above but with an additional step for the simulation of indels. We used the simulation procedure of [3], who retrieved indel-specific parameters for the rich indel model (RIM) [4] from empirical MSAs in the TreeBASE database [5]. We limited the sequence length for the indel simulation to be in $(500, 4000)$ and cropped the alignments to have a sequence length $L = 500$.

**SelReg** The SelReg model of evolution accounts for different selective regimes for each site of the alignment. It is a codon based Mutation-Selection model [6] in which each codon site is associated to one of 263 empirically assessed profiles [7] which store the fitness of each possible amino acid occurring at the site. The rate of substitution of a codon into another is stored in a $61 \times 61$ matrix and depends upon the fitness profiles, and a $4 \times 4$ matrix of mutation rates between $A, C, G, T$ [7]. Under this model, a site can evolve under neutral evolution, when all amino acids have the same fitness, under negative selection, in which case one or several amino acids have higher fitnesses and are therefore selected at the given position, or under persistent positive selection which leads the fitness profile of a site to continuously change along the phylogeny to adapt to an ever-changing selective pressure [8]. Both for training and testing we used the Pastek simulator [7] to evolve alignments along the simulated phylogenies, sampling each site's selective regime with probabilities of $25\%$, $50\%$ and $25\%$ respectively.

**Cherry** Cherry is a model of sequence evolution that represents pairwise amino-acid interactions by using a $400 \times 400$ transition matrix inferred from 15,051 Pfam MSAs with associated structure data which was used to determine contacting sites [9]. We used the Cherry model to simulate evolution along the training and testing phylogenies. The resulting MSAs, of length $L = 500$ contain $250$ pairs of adjacent coevolving sites. Although in these simulations the coevolving sites are side by side it is important to note that this does not affect in any way the presented results as none of the considered methods exploit positional information for the reconstruction of the phylogeny: the distance and ML methods assume that sites evolve independently of each other, and our network, which does not use positional encoding, is invariant to permutations of the columns in the input MSA.

##### 1.3 Detailed architecture of the Phyloformer network, training and fine-tuning

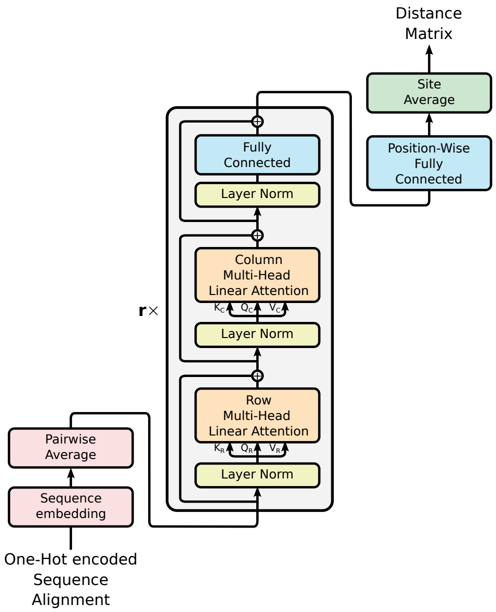

Figure 3: Network architecture of Phyloformer. We embed the one-hot encoded amino acids (a vector in $\mathbb{R}^{22}$ for each position-sequence pair in the input alignment) in $\mathbb{R}^d$ via a position-wise fully connected layer. We then take pairwise averages over sequences to obtain a representation in $\mathbb{R}^{d \times N \times L}$ of all the pairs of sequences in the alignment. Up to here each encoded amino acid is processed independently. The subsequent $r = 6$ axial attention blocks then allow them to interact building a context aware representation. After these multiple blocks we add a last position-wise fully connected layer with a softplus activation and a single output feature. Finally averaging over sites gives us the final network prediction.

The neural network architecture of the Phyloformer model is essentially that of an encoder-only transformer with Pre-Layer normalization [10, 11], with 6 attention blocks, an embedding dimension of $d = 64$, and 4 attention heads for a total number of $308,449$ trainable parameters. The network takes as input the one-hot encoded representation of an MSA as a tensor in $\mathbb{R}^{22 \times n \times L}$ and transforms it through a position-wise fully connected layer which acts identically on each amino acid representation to embed it into $\mathbb{R}^d$. We then take pairwise averages over the rows (corresponding to the sequences in the MSA) of the resulting tensor in $\mathbb{R}^{d \times n \times L}$ to obtain one in $\mathbb{R}^{d \times N \times L}$, with $N = \binom{n}{2}$, having as rows the representations of each pair of sequences from which the network will eventually predict evolutionary distances. Subsequently the tensor is processed through 6 attention blocks. We resort here to the axial attention paradigm [12], which has been employed to deal with MSA data [13], and which allows passing information across positions in two steps. First through an attention mechanism applied to each row, allowing to share information across all sites in a given pair of sequences. Second through an attention mechanism applied to each column, permitting the same information flow across all pairs of sequences at a given site. We include one LayerNorm normalization layer before and one skip connection layer after each of the row-wise, column-wise attention and fully connected layers (Fig. 3). As standard practice in the transformer literature, we use, for the final position-wise fully connected neural network inside each block, a hidden dimension $d_h = 4 \times d$, using a GELU activation function [14] following [13]. We deviate from the implementation in [13] by allowing for independent attention maps for each row, and by using the novel variant of the linear attention mechanism [15] described in section 4 of the main text. At the end of the attention blocks a last position-wise feed forward layer with a softplus activation (which ensures the positivity of the outputs) and a single output feature transforms the representation into a $\mathbb{R}^{N \times L}$ tensor, before an average over the sites gives the final network's prediction of the evolutionary distances as an $\mathbb{R}^N$ vector.

We recapitulate the training procedure for each version of Phyloformer used in our manuscript in Table 1, and the relationship between these different versions in Fig. 4.

| Network Name | Starting Network | Batch Size | Dataset Size | Model of evolution | Effective number of Steps/Epochs | GPUs used | Target learning rate | Target schedule steps | Selected checkpoint step | Loss Function |
|---|---|---|---|---|---|---|---|---|---|---|
| $\text{PF}_{\text{Base}}$ | Initialized network | 4 | 170k | LG+GC | 145.18k/20.5 | 6×A100 | $10^{-3}$ | 213.2k | 144k | MAE |
| PF | $\text{PF}_{\text{Base}}$ | 4 | 200k | LG+GC | 40.3k/4.32 | 6×A100 | $10^{-4}$ | 66k | 40,3k | MRE |
| $\text{PF}_{\text{Indel}}$ | $\text{PF}_{\text{Base}}$ | 1 | 55k | LG+GC+indels | 240k/17.45 | 4×V100 | $10^{-3}$ | 240k | 136.5k | MAE |
| $\text{PF}_{\text{Cherry}}$ | $\text{PF}_{\text{Base}}$ | 4 | 1M | Cherry | 30k/0.72 | 6×A100 | $10^{-3}$ | 66k | 18k | MAE |
| $\text{PF}_{\text{SelReg}}$ | $\text{PF}_{\text{Base}}$ | 4 | 1M | SelReg | 66k/1.58 | 6×A100 | $10^{-3}$ | 66k | 66k | MAE |

Table 1: Details of the training runs for all considered networks. We trained these networks with the Adam optimizer [16] and selected the best checkpoint based on the loss on a separate validation dataset. All models were trained using a linear learning rate schedule with 3000 steps of warmup followed by linear decay for a total of steps shown the table column. The effective number of steps the models trained for after early stopping is also shown. We either used NVIDIA A100 GPUs with 80GB of VRAM or NVIDIA V100 GPUs with 32GB of VRAM.

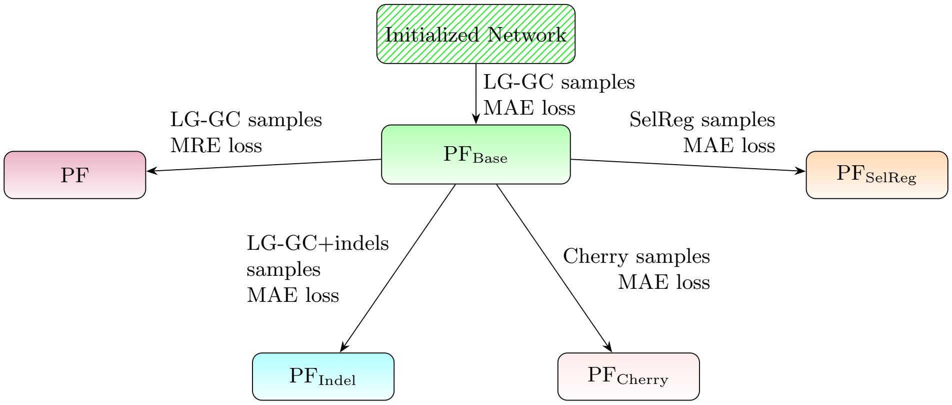

Figure 4: Trained versions of Phyloformer considered in the manuscript.

##### 1.4 Estimating posterior distributions

Here we intend to provide some intuition on why our neural network trained on simulated data provides a likelihood-free estimator of the posterior distribution $p(\tau|\text{MSA})$. More precisely for our network minimizing the MAE, it provides an estimate of the median of this distribution. We consider a two-sequence alignment for which the network outputs a single scalar predicting their evolutionary distance—and the tree $\tau$ is reduced to a single branch. Using enough sampling, we could generate several $\{(\tau_i, \text{MSA})\}_{i=1}^t$ where we sampled different values $\tau_i$ of the distance from the prior $\pi(\tau)$ which all led to sampling the *same* MSA from $p(\text{MSA}|\tau)$. The set $\{\tau_i\}_{i=1}^t$ is a sample of the unnormalized posterior distribution of $\tau|\text{MSA}$. For any particular choice of the parameters $\Phi$, our network will produce the same output $F_\Phi(\text{MSA}) = \delta$ for all these sampled alignments. Optimizing $\Phi$ over this restricted set of samples, *i.e.*, to minimize $t^{-1}\sum_{i=1}^t |\delta - \tau_i|$, would therefore lead $F_\Phi$ to output a median of the $\{\tau_i\}_{i=1}^t$, which is identical to the median of the posterior $p(\tau|\text{MSA})$.

In this artificial example, using a parameterized network would be a clear overkill, and a single scalar would suffice. In practice however, it is unlikely that we ever sample the same MSA twice or that the MSA for which we want to do inference was even once among the samples. This justifies using a parameterized $F_\Phi$ that interpolates between the $n$ available observations by minimizing $n^{-1}\sum_{i=1}^n |F_\Phi(MSA_i) - \tau_i|$ over $\Phi$. By doing so, it produces an amortized estimator for the median of the posterior distribution of $\tau|MSA$ that borrows information among close samples.

### 2 Supplementary Results

#### 2.1 Phyloformer's attention maps reveal coevolution patterns

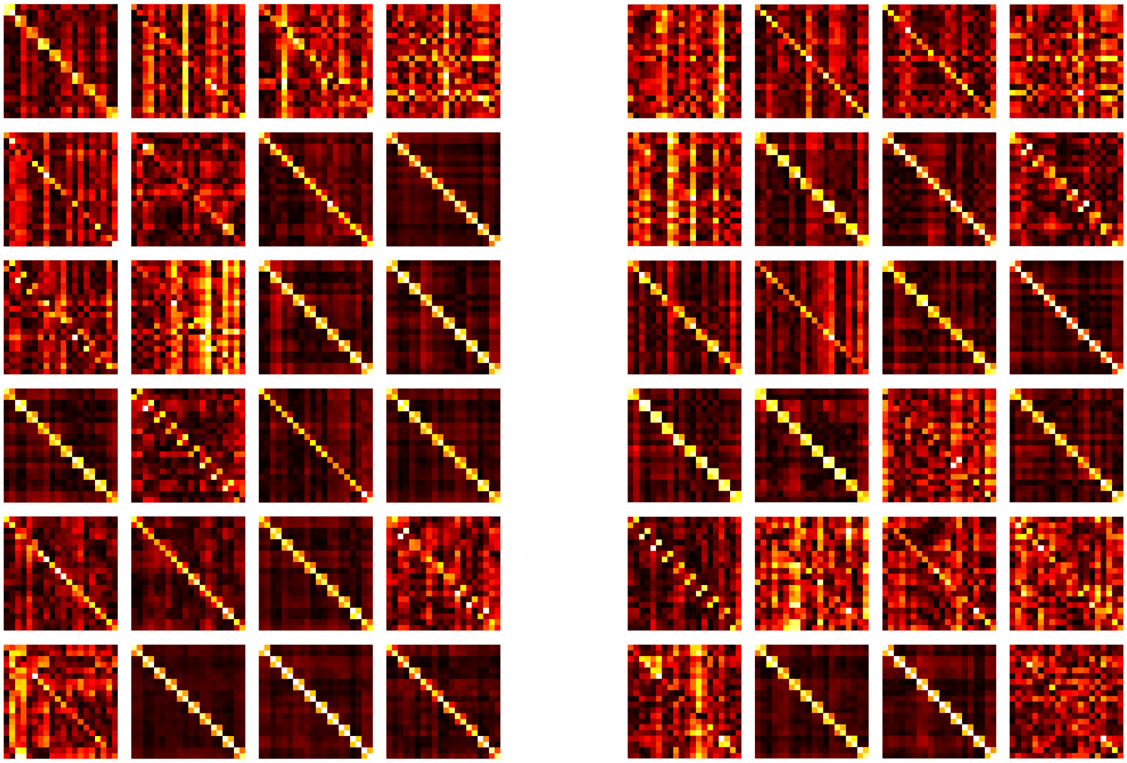

Figure 5: Row attention maps (rescaled and averaged over all alignments and sequence pairs) obtained during inference on Cherry test data with the cherry fine-tuned model (left) and the base one (right). Only the first 20 sites are shown here for ease of visualisation. Attention heads from 1 to 4 along the columns and attention blocks from 1 to 6 along the rows. The $2 \times 2$ blocks along the diagonal show that the model identifies sites that are coevolving and takes it into account for distance predictions. Interestingly the coevolution pattern can also be observed in the maps of the non fine-tuned model which has only been trained on alignments simulated without any coevolution. When the models run on the LG+GC test data without coevolution only $1 \times 1$ blocks along the diagonal can be seen (Table 2).

The linear attention approach that we use in this work does not compute attention maps explicitly. Nonetheless, we can obtain these maps by computing $A = \phi(Q) \cdot \phi(K)^T$. Focusing on attention across sites, which axial attention performs separately for each pair, we obtained $\binom{n}{2}$ maps of dimension $\mathbb{R}^{L \times L}$. Inspecting these maps can provide insight into the network's inner workings. In particular to investigate the performances of our approach in the simulations which take site coevolution into account, we computed the row attention maps (24 in total corresponding to each block and attention head) during inference on a subset (corresponding to the alignments with 20 sequences) of the test data generated both under the Cherry and LG+GC evolution model, using both the base model and the model fine-tuned on Cherry data. We averaged all $L \times L$ maps of each head and layer over all sequence pairs and further averaged over all alignments which gave us 2-dimensional matrices that can be visualized as heatmaps. To take into account both the cases where a pair of position receives particularly low or high attention score, we centered the attention maps values around 0, took their absolute value and finally normalized them to be between 0 and 1. Fig. 5 (left panel) shows that averaged attention maps obtained on data simulated with pairs of coevolving sites contain high values in $2 \times 2$ blocks along the diagonal. Remarkably we observed the same pattern on attention maps obtained on the same Cherry alignments with the network that we trained on LG+GC alignments, where each site evolved independently (Fig. 5, right panel). Our understanding is that both versions of the neural network have learned to share information between correlated sites, regardless of their position (again, no positional information is available to our network). The block pattern that we observe is just a consequence of coevolving sites being near each others in the Cherry alignments over which we compute these attention maps.

| *Network* on **test dataset**: | $\text{PF}_{\text{Cherry}}$ on **Cherry** | $\text{PF}_{\text{Base}}$ on **Cherry** | $\text{PF}_{\text{Cherry}}$ on **LG** | $\text{PF}_{\text{Base}}$ on **LG** |
|---|---|---|---|---|
| $a$, mean of $\bar{A}_{i,j}$ values, $\|i-j\|=1$ | 0.256 | 0.255 | 0.120 | 0.135 |
| $b$, mean of $\bar{A}_{i,j}$ values, $\|i-j\|>1$ | 0.098 | 0.115 | 0.121 | 0.136 |
| Ratio $a/b$ | 4.424 | 3.408 | 0.999 | 0.995 |
| mean of $\bar{A}_{i,j}$ values, $i=j$ | 0.579 | 0.535 | 0.542 | 0.523 |

Table 2: Quantitative measure of the behaviour of the base network and the fine-tuned network on the different test datasets of 20-leaves trees. We denote here by $\bar{A}$ the rescaled attention maps averaged across all 24 attention heads (4 in each of the 6 blocks) and all alignments. Whereas the values $\bar{A}_{i,j},\ i=j$, along the diagonal, are naturally higher for all combinations of network-test dataset, only when the networks are run on data containing coevolution the $\bar{A}_{i,j},\ |i-j|>1$, values stand out.

This can explain why $\text{PF}_{\text{Base}}$'s performances appear to be less affected compared to FastME and FastTree although all methods are subject to the same extent of model misspecification. Similarly this could be the reason why the base model still achieves state of the art performance across all metrics for smaller trees (Fig. 11).

On the other hand, as expected, we do not observe such pattern when running either version of the network on the LG+GC test data (Table 2) except for naturally occurring high values along the main diagonal.

#### 2.2 Phyloformer reconstructs likely trees

While working under a model amenable to likelihood calculations we can also compare the performances of the different methods in terms of the likelihood of the reconstructed tree. In Fig. 6 we can see the ratio of the log-likelihood values (computed by IQTree on the LG+GC test dataset) of the true and the predicted tree for the different methods. Higher likelihood trees lead to lower ratios. Despite the fact that our network does not explicitly optimize the tree likelihood, we observe that our method typically reconstructs trees with higher likelihoods than the true ones, as observed with the full scale ML approaches. This highlights the difference with the standard distance-based method which shows the opposite trend. Similarly when testing the different approaches on the empirical data one observe that the trees reconstructed by Phyloformer consistently have higher likelihoods than those inferred by FastME (Fig. 7, where for each tree we plot the likelihood of the one inferred by the different methods divided by that of the one inferred by IQ-TREE10 for comparison).

#### 2.3 Short branches, not short distances, explain Phyloformer's deteriorating topological performance as the number of leaves grows

Phyloformer yields larger relative errors on small pairwise distances (Figs. 17 and 19). It is natural to think that the increase in topological error along with the number of leaves might be due to a corresponding increase in smaller pairwise distances within such trees. However, Fig. 8 partly disproves this. Indeed the distribution of pairwise distances stays relatively stable across trees with different numbers of leaves with only a slight shift towards smaller distances for bigger trees. Somewhat paradoxically, the smallest pairwise distances are found in the trees with the lowest number of leaves, in particular 10-leaf trees. This is due to our simulation procedure where simulated (tree,alignments) pairs with duplicated sequences are discarded. In trees with fewer leaves, small distances do not necessarily imply small branches therefore small distances are less likely to lead to duplicated sequences than in larger trees and therefore are more likely to be part of the testing data. This important distribution shift between 10-leaf trees and trees with more leaves most likely explains Phyloformer's relatively poor performance on these small trees.

Given that the distribution of empirical tree-diameters we sample from to rescale simulated trees is the same regardless of the simulated number of leaves (see Section 1), the distribution of branch lengths within the trees changes with the number of leaves. Indeed Fig. 9a shows that the average branch length decreases as the number of leaves increases inducing a shift in the distribution of branch lengths in our simulated testing data compared to our training data. In parallel, Fig. 9b shows that Phyloformer mis-predicts shorter branches, and branches of the simulated trees that are recovered in Phyloformer trees are on average $\approx 10$ times longer than those that are not found in Phyloformer trees. Therefore, since (1) Phyloformer is more likely to make mistakes for shorter branches and (2) due to our rescaling of trees with empirical diameters, trees with more leaves are more likely to have shorter branches, it stands to reason that as the number of leaves grows so does the number of mis-predicted branches and consequently so does Phyloformer's topological error.

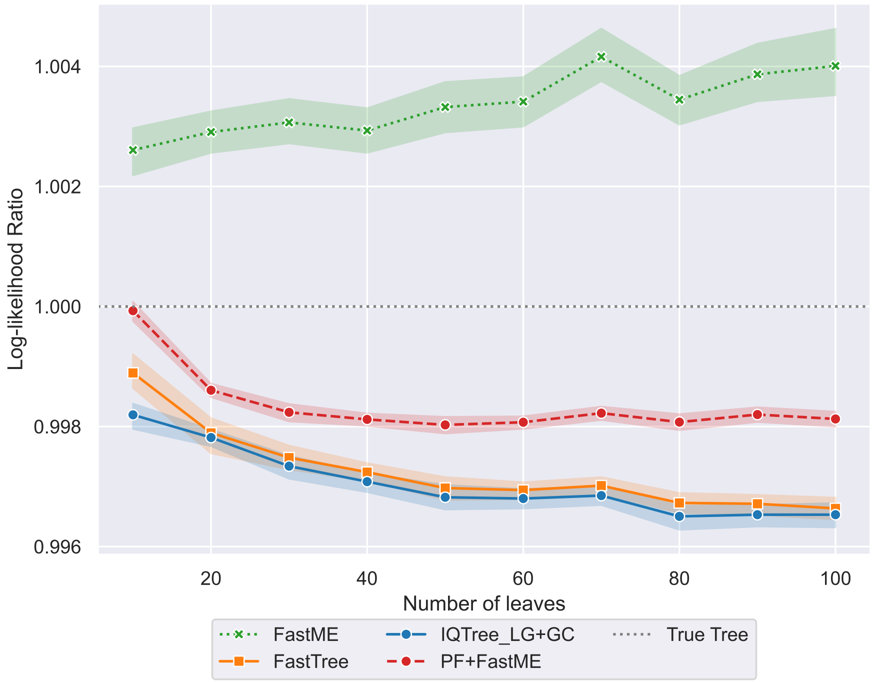

Figure 6: Log-likelihood ratios ($\mathcal{L}(T_{predicted})/\mathcal{L}(T_{real})$) for trees inferred from MSAs simulated under the LG+GC model. We use the true known simulated tree compute $\mathcal{L}(T_{real})$.

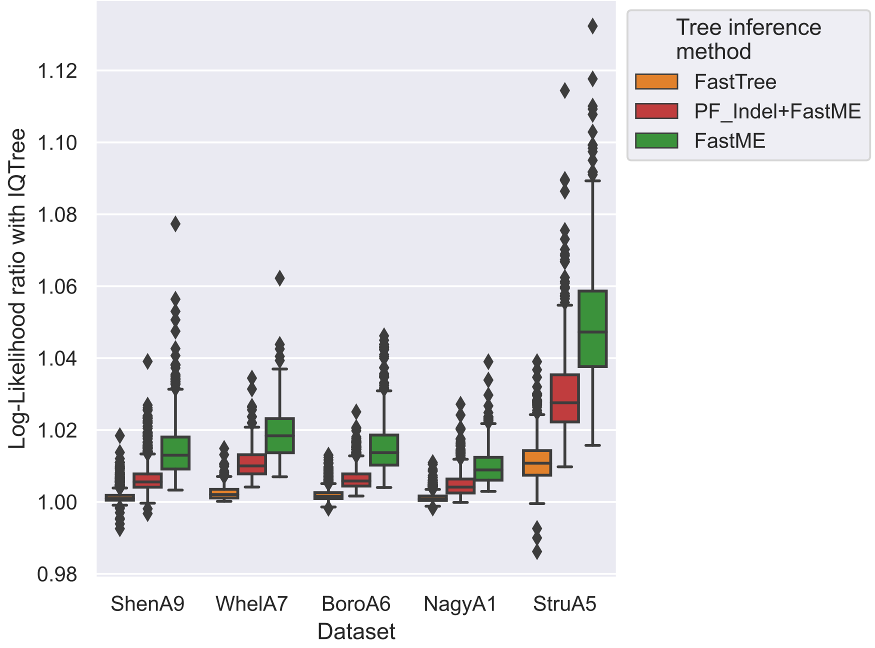

Figure 7: Log-likelihoods ratios between inferred tree and best one found by IQTree ($\mathcal{L}(T_{predicted})/\mathcal{L}(T_{IQTree})$) for the trees reconstructed on empirical alignments from [17].

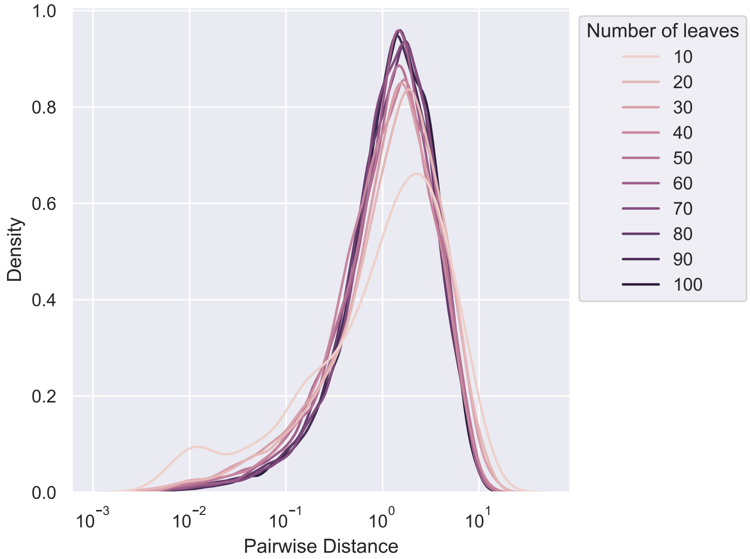

Figure 8: Distribution of pairwise distances, stratified by number of leaves in simulated testing data.

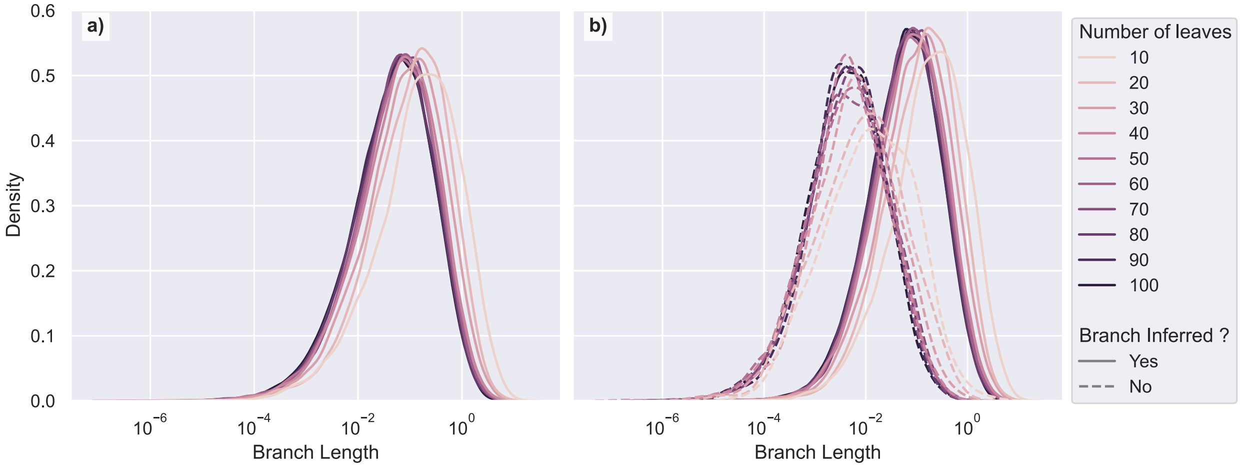

Figure 9: Distribution of branch lengths and its effect on Phyloformer performance.
**a)** Distribution of branch lengths in simulated test trees, stratified by number of leaves. **b)** Distribution of branch lengths for branches recovered or not in Phyloformer trees, stratified by number of leaves.

### 3 Supplementary Figures

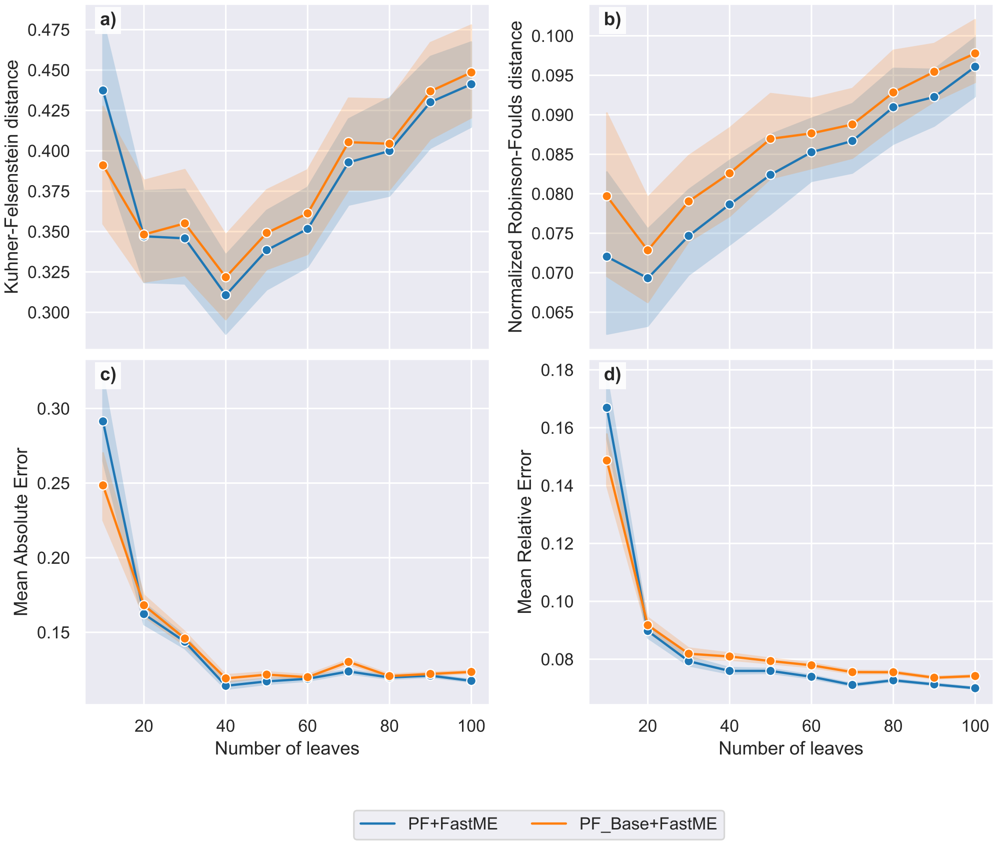

Figure 10: Performances of the base model ($PF_{Base}$+FastME) against the model finetuned with a mean relative error loss (PF+FastME). For each measure, we show 95% confidence intervals estimated with 1000 bootstrap samples.Performance is measured with: **a)** Kuhner-Felsenstein distance; **b)** normalized Robinson-Foulds distance; **c)** mean absolute error (MAE); **d)** mean relative error.

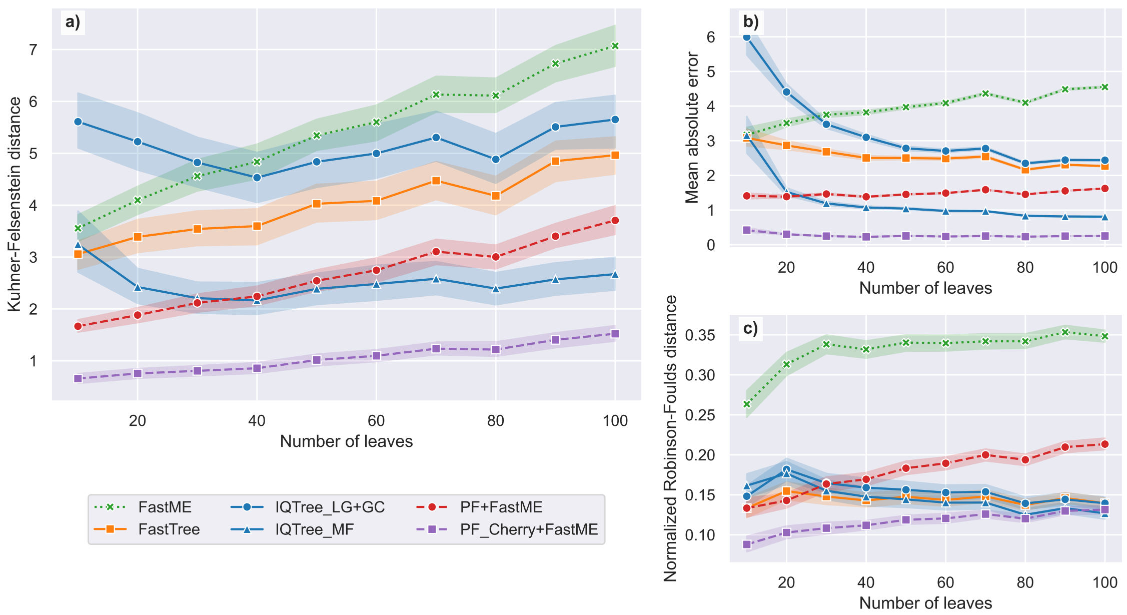

Figure 11: Performance measures for different tree reconstruction method on data simulated under the Cherry model. **a)** Kuhner-Felsenstein distance **b)** mean absolute error (MAE) on pairwise distances **c)** normalized Robinson-Foulds (RF) distance. The alignments for which trees are inferred were simulated under the Cherry sequence model and are all 500 amino acids long. For each measure, we show 95% confidence intervals estimated with 1000 bootstrap samples. Trees were inferred with (1) maximum likelihood methods: IQTree with LG+GC model, IQTree with model finder option and FastTree, (2) the commonly used FastME distance method and (3) Phyloformer models either trained on LG+GC or trained on LG+GC and fine-tuned on Cherry data.

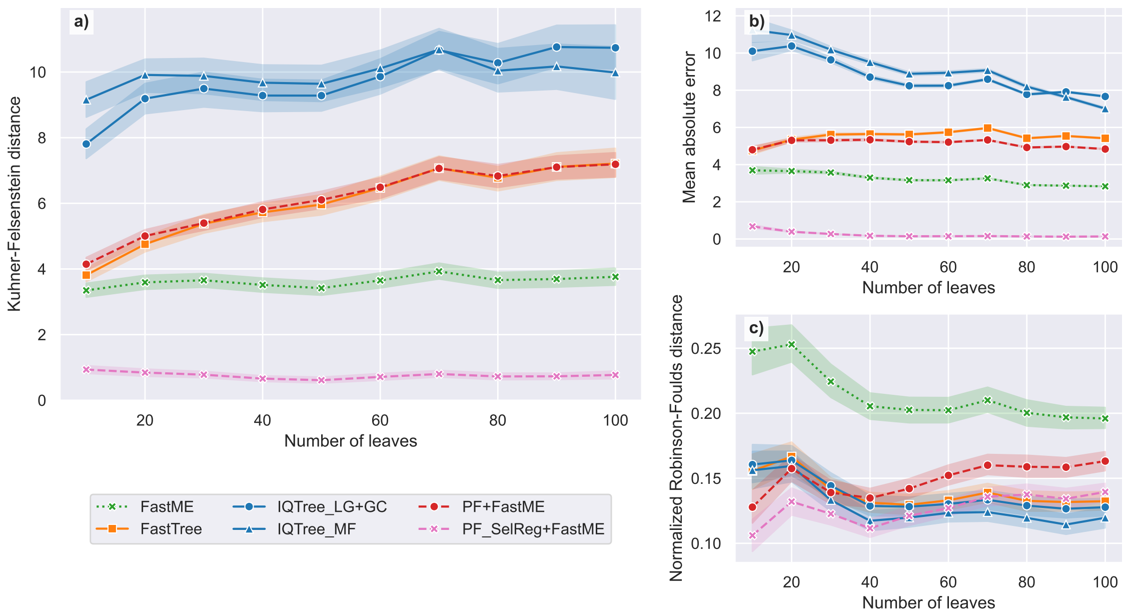

Figure 12: Performance measures for different tree reconstruction method on data simulated under the SelReg model. **a)** Kuhner-Felsenstein (KF) distance; **b)** mean absolute error (MAE) on pairwise distances; **c)** normalized Robinson-Foulds (RF) distance. The alignments for which trees are inferred were simulated under the SelReg sequence model and are all 500 amino acids long. For each measure, we show 95% confidence intervals estimated with 1000 bootstrap samples. Trees were inferred with (1) maximum likelihood methods: IQTree with LG+GC model, IQTree with model finder option and FastTree, (2) the commonly used FastME distance method and (3) Phyloformer models either trained on LG+GC or trained on LG+GC and fine-tuned on SelReg data.

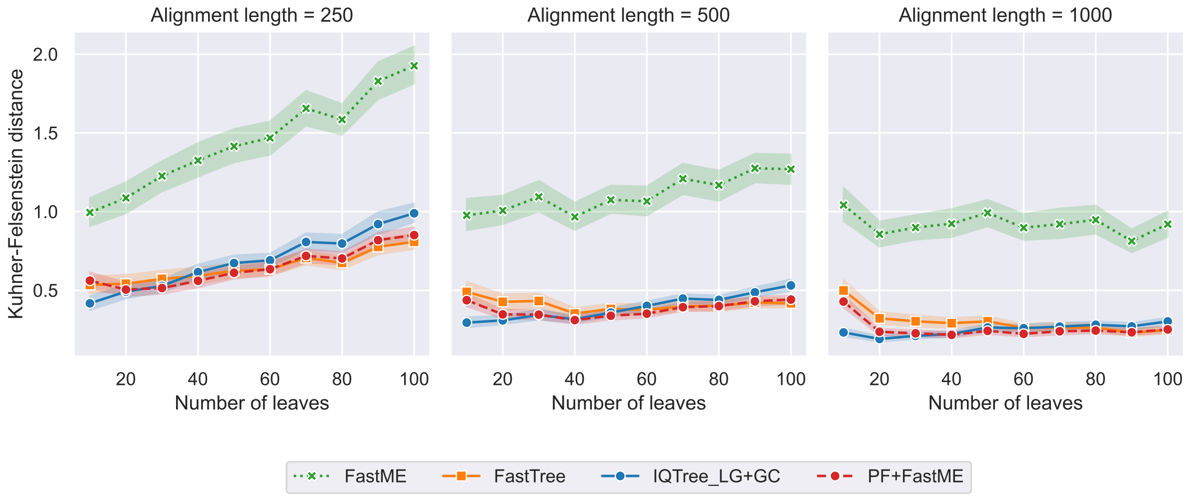

Figure 13: Effect of alignment length on the Kuhner-Felsenstein distance. Performance is measured on alignments simulated under the LG+GC model. 95% confidence intervals are estimated with 1000 boostrap samples.

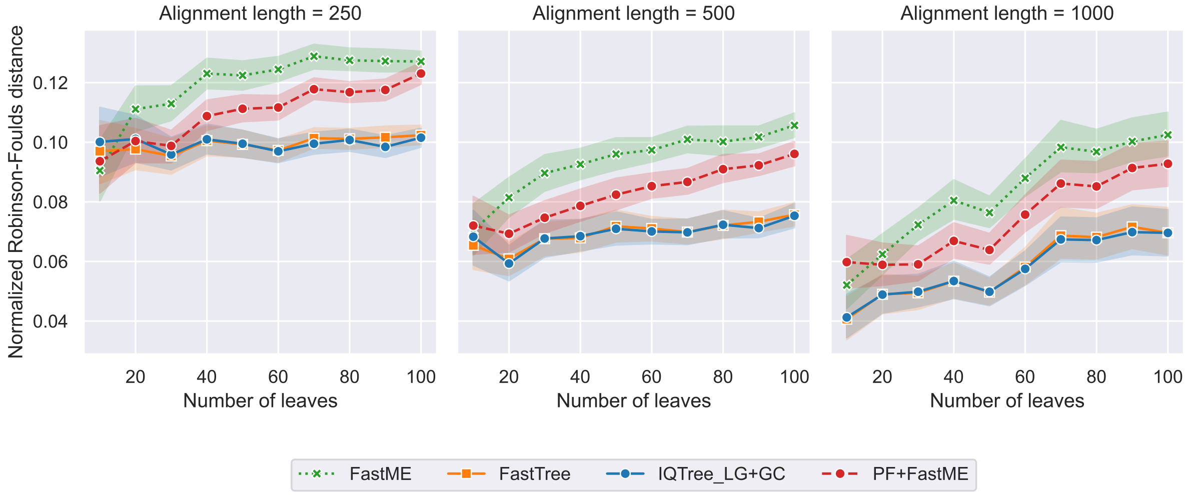

Figure 14: Effect of alignment length on the normalized Robinson-Foulds distance. Performance is measured on alignments simulated under the LG+GC model. 95% confidence intervals are estimated with 1000 boostrap samples.

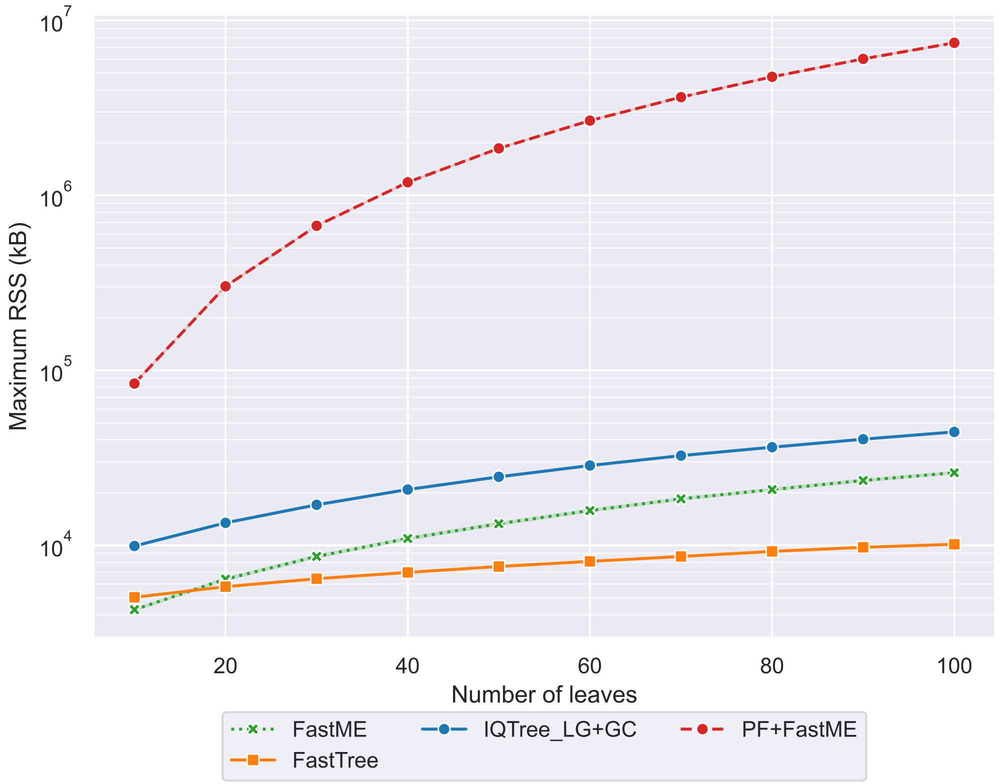

Figure 15: Memory usage during tree inference for selected tree inference methods. For methods executed on the CPU (FastME, IQTree and FastTree) the maximum resident set size (RSS), measured with the `/usr/bin/time` executable is shown. For Phyloformer, the maximum number of GPU allocated bytes added to FastME's memory usage is shown.

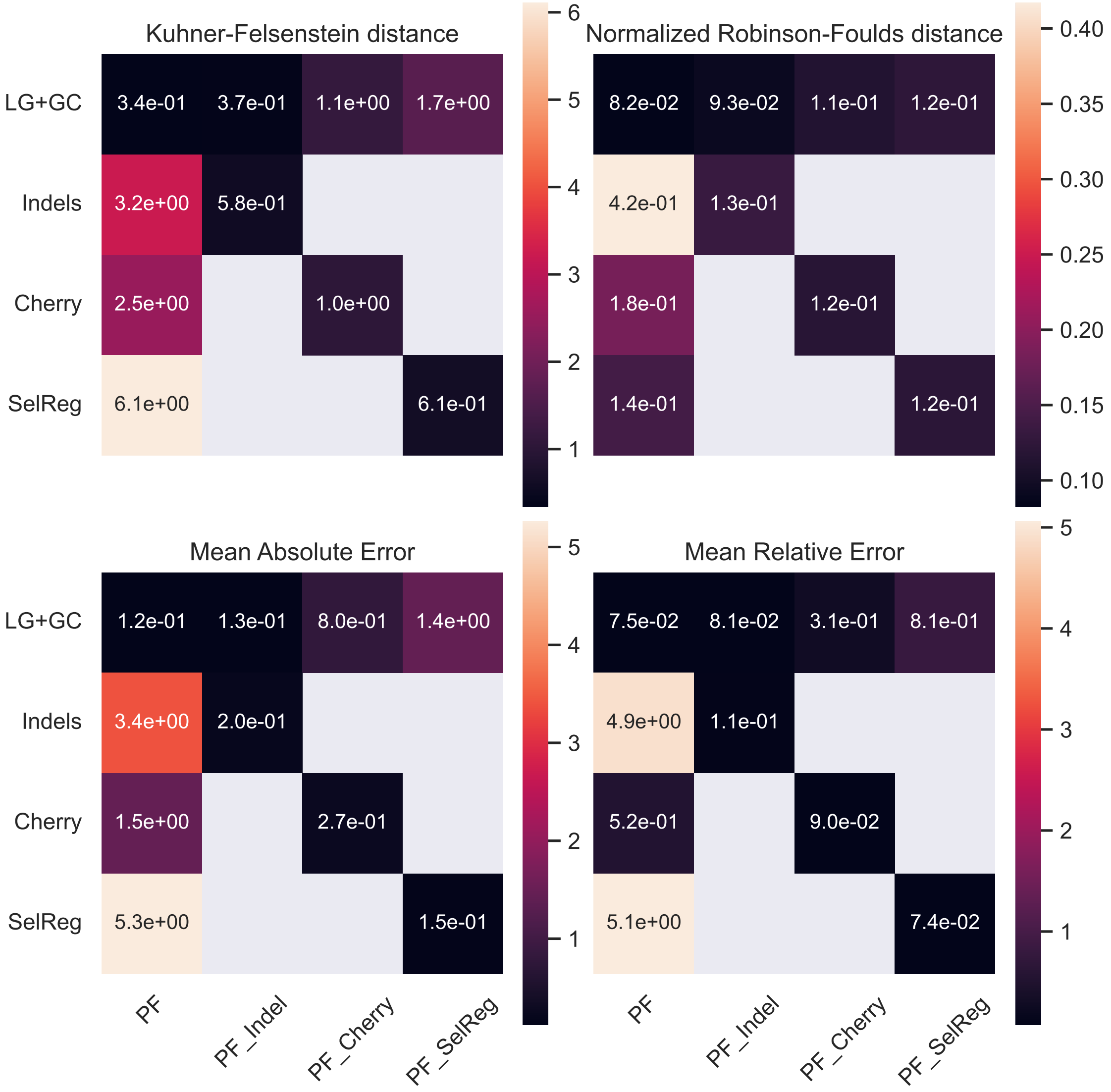

Figure 16: Effects of model specification for Phyloformer. We measured the performance of Phyloformer models (columns from left to right) fine-tuned with MRE on LG+GC, or fine tuned with MAE on LG+GC data with indels, on Cherry data or on SelReg data. For each model, the performance was measured on 50-leaf trees and corresponding alignments simulated with (rows from top to bottom) LG+GC, LG+GC with indels, Cherry and SelReg. Performance was measured with Kuhner-Felsenstein distance; normalized Robinson-Foulds distance; mean absolute error (MAE) ; mean relative error

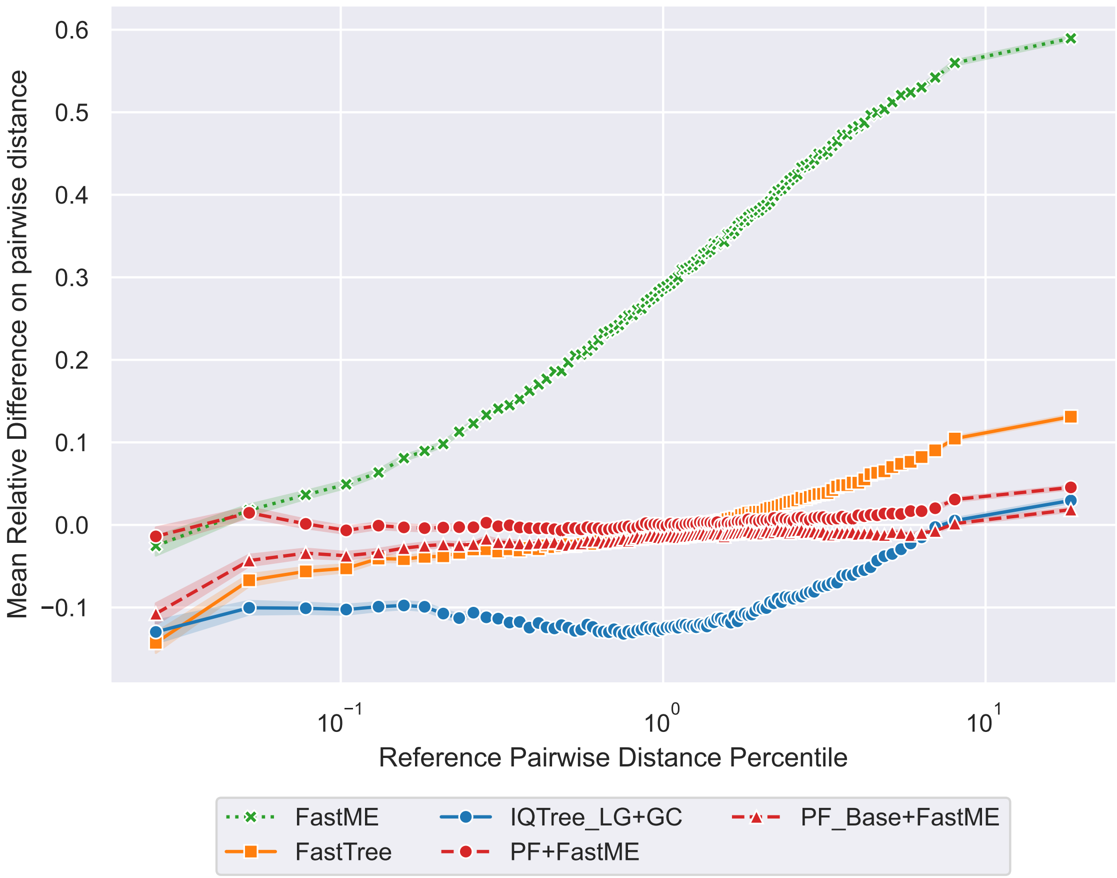

Figure 17: Mean Relative Difference (MRD) per percentile of true pairwise distances. The MRD is defined as $\ell_{MRD} = N^{-1} \sum_{i=1}^{N} d_i^{-1}(d_i - \hat{d}_i)$ with $N$ the number of pairwise distances and, $d_i$ and $\hat{d}_i$ the true and estimated pairwise distances respectively. In this figure the $\text{PF}_{\text{Base}}$+FastME model's performance is shown in addition to that of the MRE-fine-tuned PF+FastME model. The fine-tuning with MRE improved the performance on smaller pairwise distances especially with the MRD for the smallest percentile improving from $-0.1$ ($\text{PF}_{\text{Base}}$+FastME) to $\approx 0$ (PF+FastME). Overall, the curve of the MRE-fine-tuned PF+FastME is flatter and closer to $0$ than all other tree inference methods.

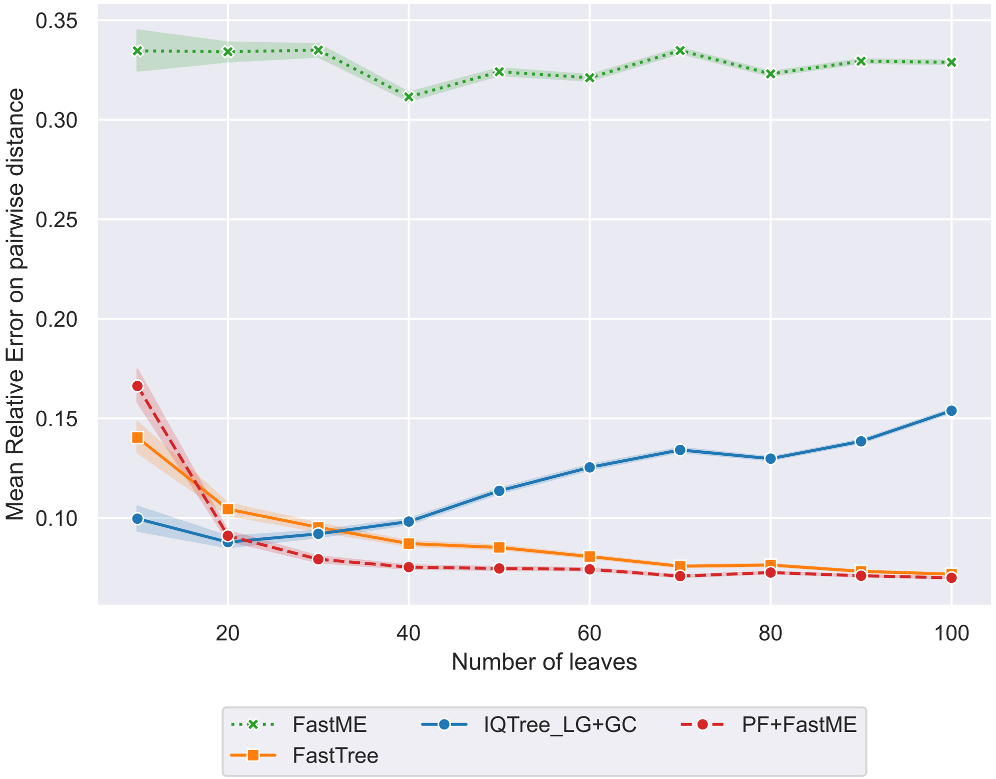

Figure 18: Mean relative error (MRE) per number of leaves for tree inference methods. Performance is measured on 500-amino acid long MSAs simulated under the LG+GC model. 95% Confidence intervals are estimated with 1000 bootstrap samples.

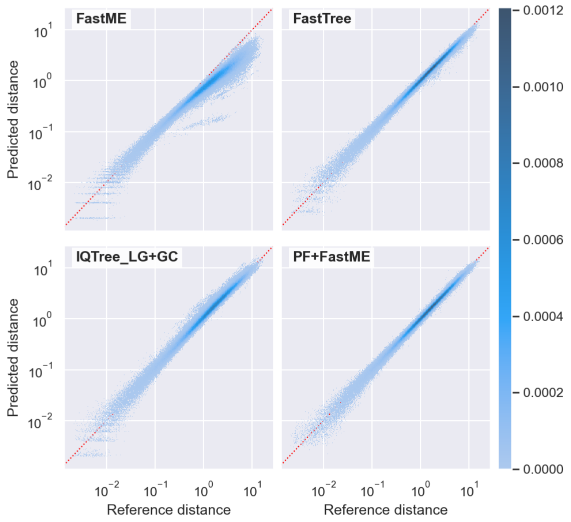

(a) LG+GC

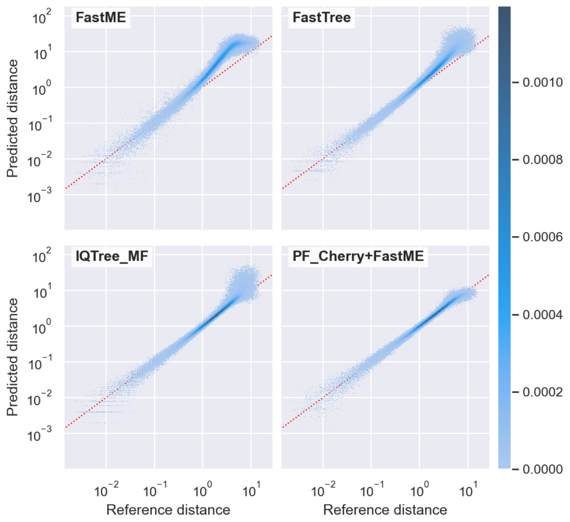

(b) Cherry

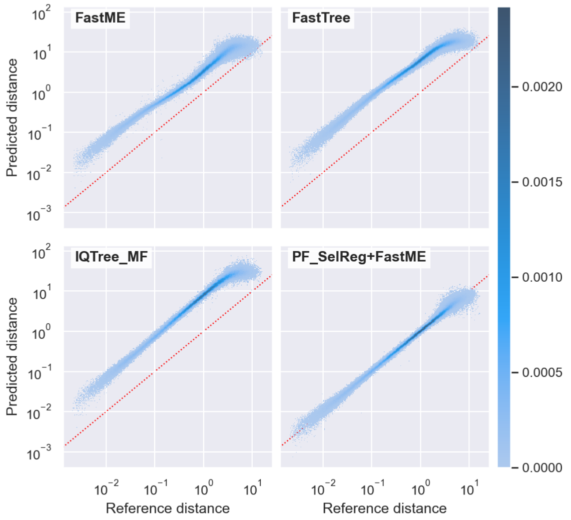

(c) SelReg

Figure 19: 2-dimensional histogram of predicted vs real distances on **(a)** the LG+GC, **(b)** Cherry and **(c)** SelReg test datasets. Under the LG+GC model (a), FastME tends to underestimate longer distances. This is expected as long distances give rise to multiple substitutions per site which are hard to detect when looking at pairs of sequences separately. PF on the other hand, exploiting the information contained in the whole alignment, recovers long distances more accurately. IQTree tends to overestimate all distances. On data simulated with Cherry (b), standard methods overestimate distances, long ones in particular. This is expected as the Cherry model considers pairs of interacting sites as states, and not the indvidual sites themselves. A substitution at the level of the pair can affect both sites, which is interpreted by site-independent models as two independent substitutions. After training on Cherry alignments, Phyloformer predicts accurate distances. On data simulated with SelReg (c), standard methods overestimate distances. Two reasons can explain this. First, the data has been simulated with site-wise vectors of amino acid equlibrium frequencies, whereas the inference model assumes that all sites share a single vector of amino acid frequencies. Second, the model of rate heterogeneity across sites used in inference (the gamma model) assumes a continuous distribution of rates, and probably does not fit well on data in which sites evolved under negative selection, neutral evolution, or positive selection. After training on SelReg data, Phyloformer predicts distances accurately.

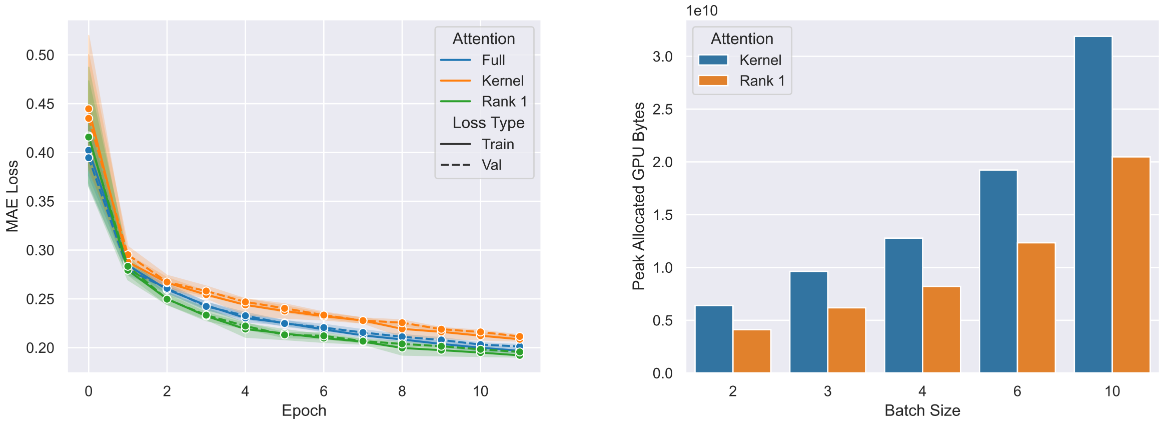

(a) MAE loss per attention function.

(b) Peak training memory usage per attention function.

Figure 20: **a)** Training and validation MAE loss per epoch for the same model with different attention functions: Full scaled dot product attention, Linear Kernel attention and our version of Rank 1 Linear Kernel attention. **b)** Peak allocated bytes on the GPU for the same model with either Linear Kernel attention or our Rank 1 version of Linear Kernel attention. All measures were done on 50 leave simulated trees.

[10] Ashish Vaswani, Noam Shazeer, Niki Parmar, Jakob Uszkoreit, Llion Jones, Aidan N Gomez, Łukasz Kaiser, and Illia Polosukhin. Attention is all you need. *Advances in neural information processing systems*, 30, 2017.

[11] Ruibin Xiong, Yunchang Yang, Di He, Kai Zheng, Shuxin Zheng, Chen Xing, Huishuai Zhang, Yanyan Lan, Liwei Wang, and Tieyan Liu. On layer normalization in the transformer architecture. In *International Conference on Machine Learning*, pages 10524–10533. PMLR, 2020.

[12] Jonathan Ho, Nal Kalchbrenner, Dirk Weissenborn, and Tim Salimans. Axial attention in multidimensional transformers. *CoRR*, abs/1912.12180, 2019. URL `http://arxiv.org/abs/1912.12180`.

[13] Roshan M Rao, Jason Liu, Robert Verkuil, Joshua Meier, John Canny, Pieter Abbeel, Tom Sercu, and Alexander Rives. Msa transformer. In Marina Meila and Tong Zhang, editors, *Proceedings of the 38th International Conference on Machine Learning*, volume 139 of *Proceedings of Machine Learning Research*, pages 8844–8856. PMLR, 18–24 Jul 2021. URL `https://proceedings.mlr.press/v139/rao21a.html`.

[14] Dan Hendrycks and Kevin Gimpel. Gaussian error linear units (gelus). *arXiv preprint arXiv:1606.08415*, 2016.

[15] Angelos Katharopoulos, Apoorv Vyas, Nikolaos Pappas, and François Fleuret. Transformers are rnns: Fast autoregressive transformers with linear attention, 2020.

[16] Diederik P. Kingma and Jimmy Ba. Adam: A method for stochastic optimization, 2015. URL `http://arxiv.org/abs/1412.6980`.

[17] Xiaofan Zhou, Xing-Xing Shen, Chris Todd Hittinger, and Antonis Rokas. Evaluating fast maximum likelihood-based phylogenetic programs using empirical phylogenomic data sets. 35(2):486–503. ISSN 0737-4038. doi: 10.1093/molbev/msx302. URL `https://doi.org/10.1093/molbev/msx302`.
